# Silk Road Revealed: Mechanism of silk fibre formation in *Bombyx mori*

**DOI:** 10.1101/2023.06.02.543394

**Authors:** R.O. Moreno-Tortolero, Y. Luo, F. Parmeggiani, N. Skaer, R. Walker, L. Serpell, C. Holland, S.A. Davis

## Abstract

The transition of silk fibroin from liquid to solid is fundamental to silk-fibre production and key to the superior materials properties of native silks. Here we discover that the fibroin heavy chain from the silkworm moth *Bombyx mori* folds into a novel β-solenoid structure, where the N-terminal domain (NTD) promotes higher-order oligomerization driven by pH reduction. These findings elucidate the complex rheological behaviour of silk and the liquid crystalline textures within the silk gland. We also find that NTD undergoes hydrolysis during standard regeneration, explaining differences between native and regenerated silk feedstocks. Overall, this study establishes a fibroin heavy chain fold, which could be relevant for other similar proteins, and explains mechanistically its liquid-to-solid transition, driven by pH reduction and stress.

**One sentence summary:** This study redefines the molecular structure of fibroin heavy chain and its role in the transition from solution to fibre.

## Main Text

Within the larval insect the major protein component of silk fibres, fibroin, is stored as a liquid, sustaining a triggered conversion to a very stable solid material, the silk fibre. This transformation starts within the silk gland, a highly specialised sack-shaped organ where not only the silk proteins are produced but where tight control over pH and metal ion concentrations is exerted (*1*). However, the mechanism of this transformation at the molecular level remains largely unknown. There are some commonly agreed principles, including the contribution of pH and shear stress/strain, energy input/work (*2*), in triggering assembly and promoting structural conformational changes (*3, 4*). Though, there is less consensus regarding the conformational changes at the molecular level. In particular, the molecular structure of the liquid state silk, often called silk-I, is heavily debated (*5*–*9*) and is the focus of our research.

Fibroin, both within the gland and in the fibre, is composed of three different proteins, fibroin heavy chain (FibH), fibroin light chain (FibL) and a glycoprotein, fibrohexamerin (P25) of molecular masses 392, 28 and 25 KDa, respectively. These form a complex known as the elementary unit with molar ratios of 6:6:1, wherein FibH and FibL form a heterodimer stabilized by a single disulphide linkage located at the C-terminal domain of FibH and the interaction between these and P25 is non-covalent (*10, 11*). However, the structural details of this system are not well known, except for the crystal structure of the N-terminal domain (NTD) of FibH. It is generally accepted that only FibH is essential for fibre formation, with FibL and P25 being auxiliary in the secretion process (*12*–*14*) with some silk-producing moths and other related species lacking these latter proteins entirely (*15*–*17*). Accordingly, due to its dominant mass contribution, FibH is thought to be the main protein responsible for the properties of fibroin across different length-scales, from the NMR chemical shifts (*18*), to the diffraction patterns of Silk-I (liquid state) and Silk-II (fibre) (*19*).

Silk-I has previously been termed α-silk, as it was erroneously assumed to include α-helical folds, to distinguish between this conformational polymorph and the better characterised Silk-II, which is known to be rich in antiparallel β-sheet structure (*20*). The exact structure of silk-1 remains ambiguous, and it is often classified as either a random coil or an intrinsically disordered state. However, discrete reflections in X-ray diffraction data suggest neither of these classifications is fully adequate (14,15). Currently, the most accepted Silk I model is a type-II β-turn rich structure (*23*). However, this model requires extensive intermolecular hydrogen bonding networks to persist (*24*), failing to account for the observation of similar chemical shifts both in dilute solutions and in the solid state measured by NMR, as well as small fibrillar structures observed herein and previously (*25*) by electron microscopy. Similarly, modelling of consensus sequences has indicated a right-handed β-helix showing lower relative free-energy values than the corresponding type-II β-turn (*24*). Moreover, the observation of a range of distinct liquid crystalline optical textures *in vivo* between the anterior section of the middle gland and the spinneret (*26*) remains unexplained, with the mesogenic structures being unidentified. Silk goes from an isotropic texture in the posterior and middle gland sections to a series of complex transitioning optical textures. At the start of the anterior section of the silk gland, a cellular optical texture is observed, which transforms to an isotropic phase prior to reverting to a fully nematic phase before the spinneret (*27*). The emergence of the cellular optical texture has been attributed to epitaxial anchoring of rod-like mesogens under confinement (*28*). However, this model does not adequately account for the subsequent transition to a nematic texture, under flow, as the tube diameter in the gland decreases towards the spinneret. Curiously, at a similar position to the cellular optical texture, evidence of cholesteric order has been observed using electron microscopy (*29*). More importantly, despite the evidence of supramolecular order, a transition from the Silk I to Silk II polymorph only occurs later near the spinneret (*27, 29*).

In this study, we propose that at the molecular level, Silk-I corresponds to a folded fibrillar conformation of FibH. Furthermore, we rationalise how physicochemical triggers such as pH, metal ions, and stress/strain induce conformational changes in this mesogen that result in concomitant supramolecular assembly or reorganization consistent with the observed optical textures inside the silk gland. The abrupt loss of the cellular texture derives from disruption of the preceding cholesteric phase, and is a key intermediate to facilitate the minimum energy transition to the ordered nematic phase, as previously proposed (*27, 29*). To investigate this hypothesis, we employed a comprehensive approach combining both computational simulations and experimental methods, encompassing various length scales ranging from the molecular to macroscopic. Our methodology involved the utilization of a fibroin solution achieved by dissolving silk fibres, after a proprietary degumming method, in concentrated LiBr solutions, resulting in a material referred to as native-like silk fibroin (NLSF). This NLSF exhibits numerous characteristics consistent with those of the native silk fibroin (NSF), including electrophoretic mobility and rheological properties. It is important to differentiate NLSF from standard regenerated silk fibroin (RSF) (*30*), as the latter fails to accurately replicate the rheological and mass properties of the native system.

### Molecular model creation and validation

Dilute solutions of NLSF were cast and left to dry slowly prior to being characterised by fibre X-ray diffraction (fXRD) experiments. When the incidence of the X-ray beam was perpendicular to the film, we obtained a pattern similar to those obtained by powder X-ray diffraction (pXRD), yet, as we rotated the film, sharper reflections were observed and eventually a highly oriented pattern emerged (Fig. 1A and fig. S1), consistent with previous Silk-I data (*22, 31*), see Table S1 for reflections. This data coupled with the observation of liquid crystalline phases suggest the presence of rod-like molecules, which would be preferentially oriented parallel to the surface of the film (*32*). After running Alphafold2 (AF2) simulation of the repetitive domains of FibH (*33*–*35*), we found that these repetitive domains are predicted to adopt a novel β-solenoid conformation (fig. S2). Briefly, the fold results from a supersecondary folding of strands, stabilised by inter-strand hydrogen bonds running along the solenoid axis, as depicted in Fig. 1B. Although in this case the abundance of glycine and the absence of a tightly packed core lead to motifs structurally closer to a polyproline-II or polyglycine-II configuration, which can be frequently assigned as disordered by CD spectroscopy (*36*). It is possible that other G-rich proteins, such as fibroin heavy chain-like, spidroins, and others with hexapeptide repeated motifs of GX fold in similar structures (*37*). Structural evidence of different spidroins folding into flexible worm-like structures was recently found by SAXS (*38*), pointing at the possibility that β-solenoids might be widespread among silk proteins. Comparable conformation and interpretation were found for the G-rich snow flea antifreeze protein, which was crystallized using racemates (*39, 40*). Similar β-helical configurations have been suggested before for FibH, mostly via computational approaches (8, 24).

**Fig. 1.**
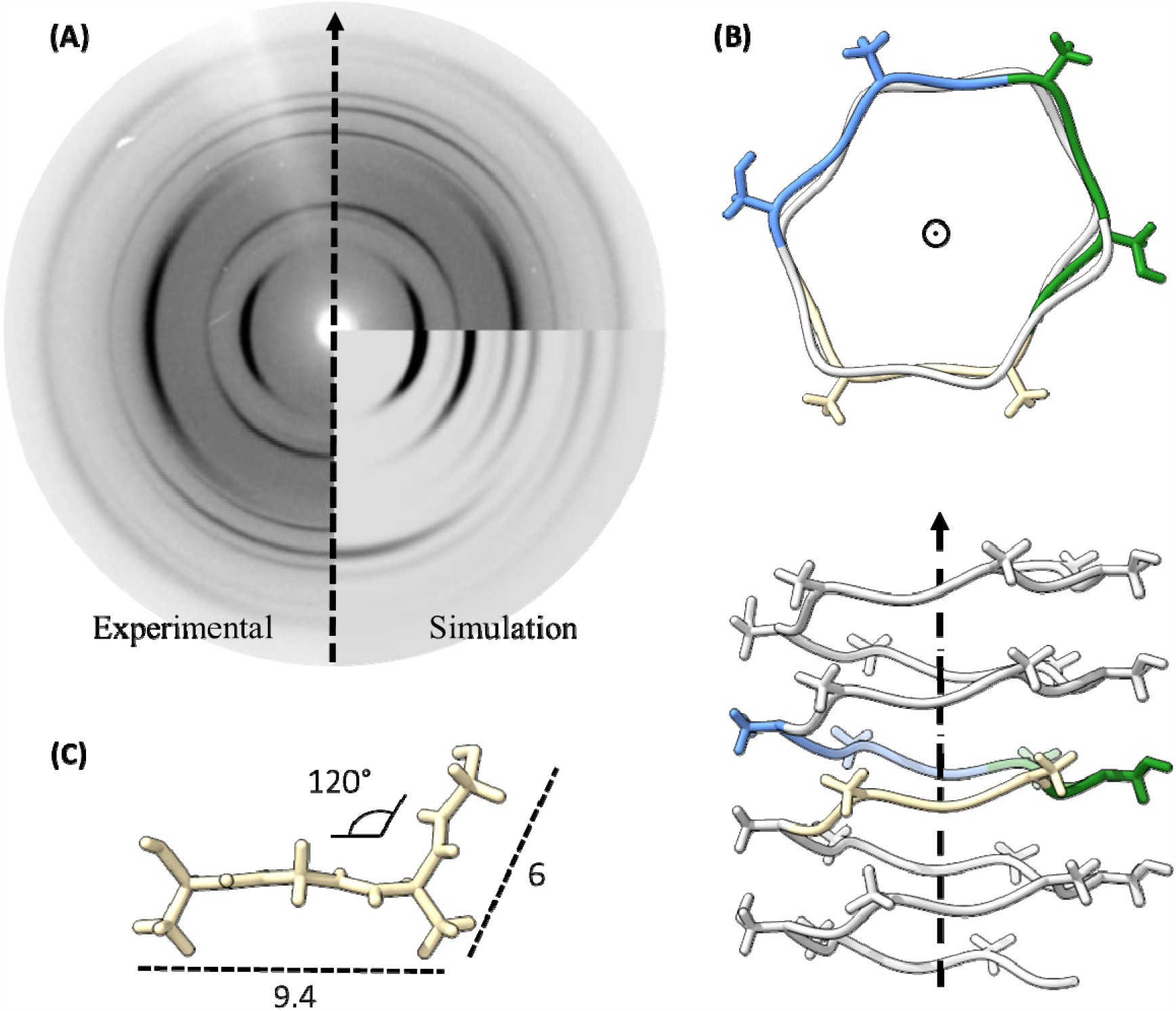
Structural model for fibroin heavy chain. (A) Experimental and simulated Silk-I diffraction pattern obtained from drop-casted film and derived unit cell showing the proposed fibre axis. (B) Proposed unit cell comprised of AGXG motifs. (C) β-solenoid molecular model obtained for fibroin heavy chain, observed from the top and side views (top and bottom, respectively).

Although the predictions showed low confidence, and relatively high predicted Local Distance Difference Test (pLDDT) with an average of about 50, all generated models maintained the solenoidal topology (fig. S3) and were consistent with the topologies we observed by TEM (fig. S4). Based on these results, we propose a trigonal unit cell containing a single curved strand, as depicted in Fig. 1C; unit cell parameters are a: 9.4, b: 3.4 and c: 6 Å. We note that the proposed unit cell is similar to that proposed for the PolyGly-II polymorph (*41*), known to resemble Silk-I (*23*) and gives a simulated diffraction pattern that recapitulated the reflections obtained experimentally (Fig. 1A). Nevertheless, observed discrepancies can be attributed to experimental constraints in aligning the sample, inherent heterogeneities in a real sample, and the software’s inability to account for helical symmetry. This unit cell is shown in context of the proposed solenoidal model in Fig. 1B. Given the markedly repetitive characteristics of FibH, featuring extensive low-complexity domains, characterized by a hexapeptide sequence (GAGAGX), where X may assume any of the residues A, S, Y, or V in descending order of abundance, twelve solenoidal domain are expected to form. The projected architecture of the protein consists of 12 solenoid bodies, interconnected by short linker regions that appear structurally unorganized, as depicted in fig. S5.

Furthermore, both 1D proton (fig. S6) and 2D ^1^H-^1^H TOCSY (fig. S7), ^1^H-^13^C HSQC, ^1^H-^15^N HSQC (fig. S8), ^1^H-^15^N TROSY (Fig. 2A) and ^1^H-^1^H NOESY (Fig. 2A and B) were acquired and assigned chemical (Table S2) shifts were very close to those reported elsewhere (*42, 43*), and predicted shifts from the solenoid models (Table S3). Notably, we did not observe changes in the chemical shifts at pH 8 or pH 6, intended to replicate physiological pH change within the silk gland (fig. S9). The assignment of the amide protons and nitrogen chemical shifts is depicted in Fig. 2A for simplified motifs within FibH. In addition, we note that when predicting the torsion angles for reduced motifs in FibH, two populations for G residues are possible within the generic GX motif (*18*). One of these (□, φ = 77°, 10°) was used to derive the type-II β-turn model for Silk-I. However, the second population of torsion angles (ca. □, φ = -80°, 180°) was not used to constrain the modelling and matches closely those found in the solenoidal model predicted by AF2 as well as previously proposed angles from density functional theory (DFT) calculations on NMR values (*9, 44*). This conformation implies that in the transformation from Silk-I to Silk-II, G residues would not need to go through sterically hindered angles (see fig. S10) (*45*), thus reducing the overall energetic cost of the Silk-I to Silk-II transformation. Further to these observations, NOESY experiments, shown in Fig. 2B evidences Y residues close to (∼ 3.3 to 3.4 Å) the methyl sidechain of A, (fig. S11), suggesting a possible methyl-pi interaction (*46*), which could be an inter-strand (within the solenoid) stabilising interaction. There are numerous Y residues on the predicted models, with the aromatic group hovering over methyl groups from A residues on contiguous strands (see Fig. 2B, bottom). Similarly, we observed other NOE intensities, at the amide proton from S residues, which although showing the strongest correlation with G, it also showed correlation with A-βH. Such patterns can be readily explained by our model, where the strongest correlation corresponds to abundant motifs gS (ca. 2.2 Å), and the weaker correlation with A-βH corresponding to across strand gS--A distance (2.7 Å), both the estimated distances and ratios are in good agreement with the experimental data (more in Supplementary text: NOESY).

**Fig. 2.**
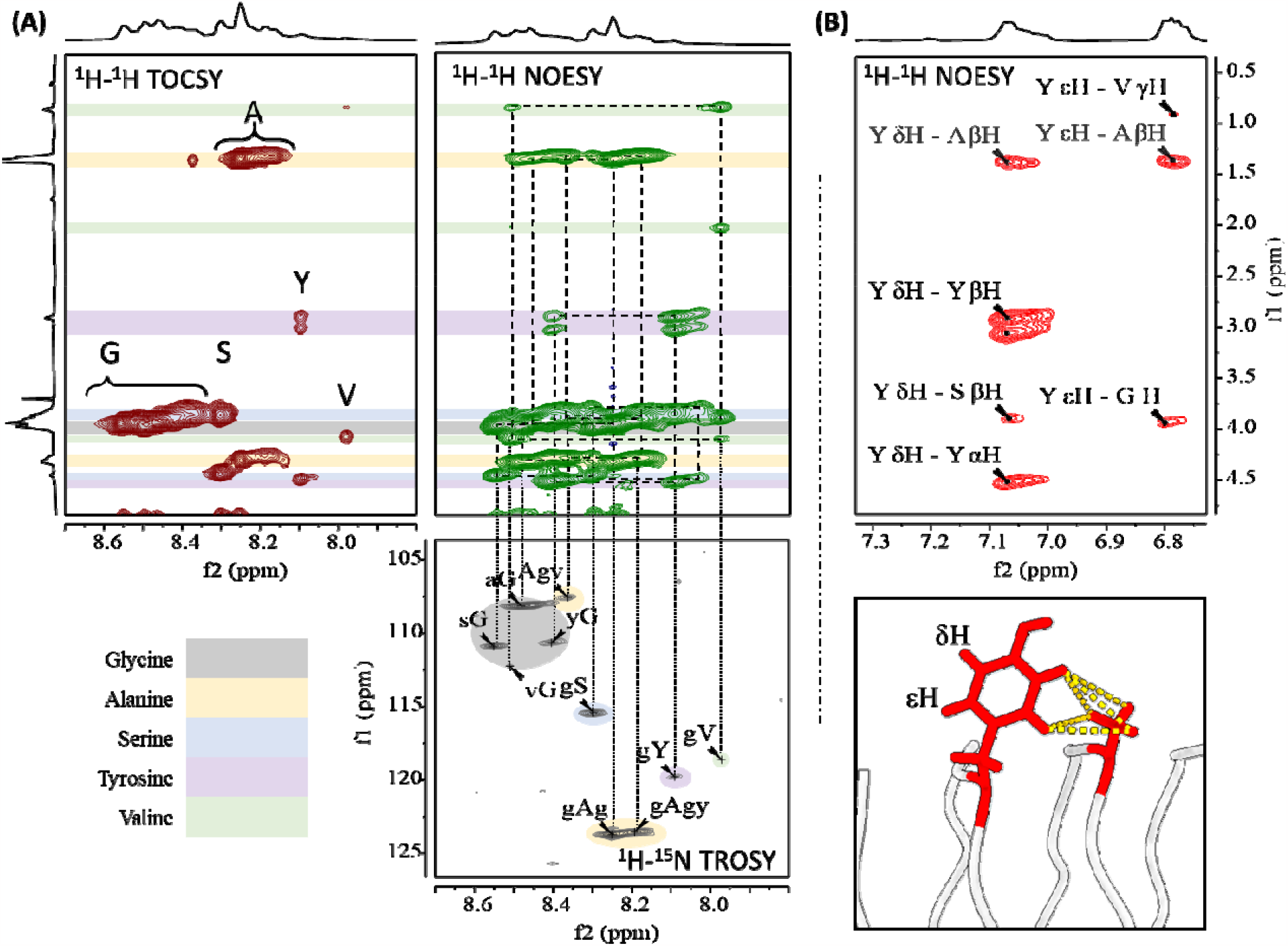
NMR analysis of native-like silk fibroin: (A) Amide shift assignment via 1H,1H TOCSY, Amide region 1H,1H NOESY analysis, and 15N chemical shift assignment of simplified motifs via 1H, 15N TROSY. (B) Analysis of the Tyrosine (Y) residue NOE signals showing evidence of proximity to Alanie-CH, and cartoon representation of motif found within the Alphafold2 model showing positions like the measured via NOESY experiment.

### The role of NTD, pH effect on rheological properties

It has largely been recognised that in native spinning both shear stress and pH changes are responsible for the transformation of the soluble fibroin into the solid fibre (*3, 47*). Rheological characterisation of native feedstock has provided snapshots into the strain-induced assembly. In most studies the silk dope is extracted from the posterior segment of the middle gland, where the pH is near neutral (*48, 49*). However, direct evidence of the structural changes induced by pH and their effects on rheological properties has proved elusive.

Here we used NLSF samples obtained at concentrations of 60 to 80 mg/mL, lower than those found inside the gland (190-300 mg/mL) (*49*), buffered at pH values to replicate the pH gradient within the silk gland, going from pH 8 in the posterior section down to 6.2 in the anterior section (*50*). We observed that the material goes through a reversible sol-to-gel transition when buffered above pH 7, with the sample at pH 7 separating into two clear phases, gel-like and liquid-like. Single phases were observed for samples either below or above pH 7. Using NLSF at pH 7, under oscillatory rheology testing we were able to replicate many previous features seen in NSF (i.e. observation of crossover), with the exception of a lower overall modulus and viscosity and a lack of a single relaxation mode in the terminal region, both of which we attribute to a lower concentration and increased polydispersity of the NLSF against NSF studies (*49*). Furthermore, we were able to uncover a reversible pH driven sol-gel transition occurring at a critical pH of 7, similar to the pH-dependent oligomerisation of NTD (*51*). Fig. 3A shows two master curves demonstrating the effect of pH on the obtained material; master curves were created from the data shown in fig. S12 and the applied shifts are presented in Table S4; the lineal viscoelastic limit was determined beforehand, and plots are shown in fig S13. Decreasing the pH further turns the gel opaque irreversibly, providing evidence of larger aggregates forming. However, no evidence suggests that the pH drops below 6 within the gland.

**Fig. 3.**
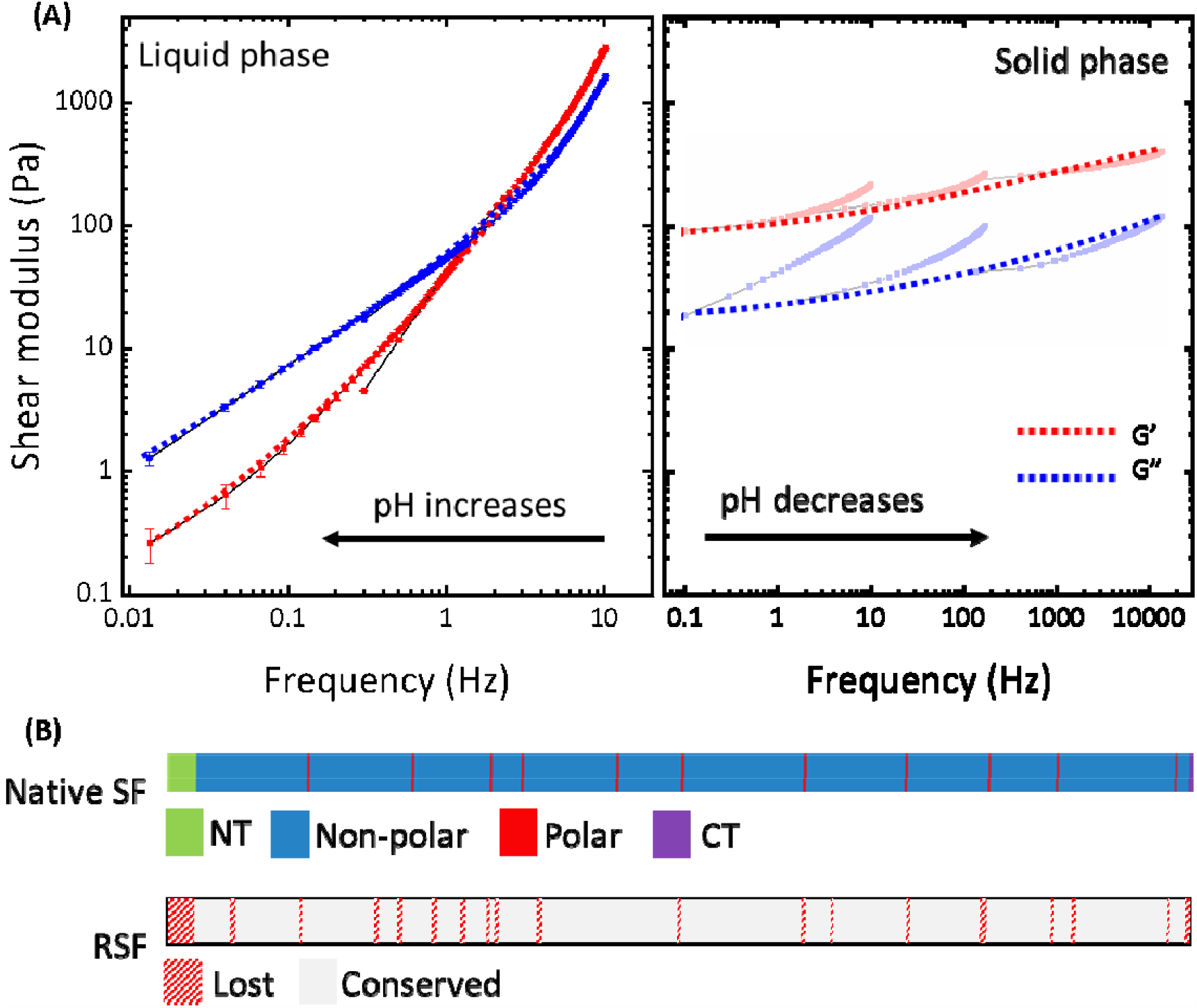
The effect of pH on rheological properties of silk fibroin, and the lost domains during regeneration. (A) Master curves created using oscillatory shear data of the liquid and solid-like samples shown on the left and right, respectively. (B) Carton showing the multidomain architecture of the primary structure of FibH (top), followed by an aligned cartoon representing the total sequence coverage from the hydrolysate peptides recovered from RSF.

For samples below pH 7, the observed shear modulus showed a reciprocal relationship with pH, increasing as the pH decreased. This suggests a more stable molecular network is formed as the pH decreases, evidenced by increased elastic modulus. Crucially, NLSF at pH 7 (liquid-like) showed curves like those reported for the extracted liquid SF from the gland, suggested to be “aquamelts,” due to their similarity to typical Maxwellian polymer melts (*52*). Notably, the cross-over frequency seemed to be shifted towards higher frequencies as pH increased, indicating a shortening in the lifespan of the interaction. Again, at pH lower than 7, the observation of gel phases indicates long-lived and increased stability of the formed structures. In all cases, given that the lineal viscoelastic limit (LVE) of the material was not exceeded, samples remained clear, and repeated cycles showed similar behaviour.

Similarly, when measuring viscosity against shear rate for samples at pH 7, the material undergoes shear-thinning until the shear viscosity increases, preceded by a normal force increase, resulting in fibrous aggregates akin to NSF (fig. S14, *53*). Interestingly, under similar experimental conditions, this transition seems to be hindered at higher pH values (something also seen for native silks and our samples in LiBr (*54*), in congruence with the dominant liquid character and the decreased stability of the interaction that gives rise to the slowest relaxation mode (*55*).

The proximity of the critical pH observed here with that reported for NTD (below pH 7, *51*), strongly hints at this domain being involved in the observed structural transitions. Furthermore, the transition is also reflected in an earlier onset of aggregation observed at pH lower than 7 when temperature ramps are conducted on CD and DLS, figs. S15-17. Consequently, we hypothesise that the absence of NTD would render the material pH-insensitive within the range (pH 8-6) and such a rheological transition would not be seen.

As a means of comparison and in support of our use of NLSF as a model system for NSF, RSF was also studied for its rheological behaviour and response to pH changes. It is well known that silk fibroin behaves very differently after standard regeneration/reconstitution (*56, 57*). For example, when shear is applied to NSF extracted directly from the posterior part of the middle gland, higher-order assembly into nanofibrils is observed, contrary to RSF at a similar pH of 7 (*3, 58, 59*). Similarly, reversible sol-gel transitions have been observed only for NSF when changing pH (*47*), while our RSF is insensitive to pH in the studied range. It is possible to promote assembly into nanofibrillar structures in RSF, but only when the pH is decreased to near the theoretical isoelectric point (PI) of FibH (ca. 4.7) (*60*), well below the physiological pH of the gland (*56, 58, 61*).

Many authors argue that the molecular weight (MW) reduction in RSF is solely responsible for these observations (62). However, we considered this reduction insufficient to explain the significant differences in literature found viscosities, or the complete insensitivity to pH (between pH 8 and 6) that we observed. Reported viscosities obtained from NSF are far superior to those reported for RSF even at similar weighted concentrations (between 500-4000 Pa•s for SF and between 0.1-1 Pa•s RSF) (49, 63). Such a significant difference, ∼4 orders of magnitude, is not substantiated by the MW difference, which on average only falls by approximately half after 30 mins of boiling during degumming (64), as depicted in fig. S18. This reduction in MW would be translated in a proportional reduction of ∼0.088 (calculation in Supplementary text: polymer viscosity), close to one order of magnitude drop in viscosity (3 orders of magnitude away from the experimental difference). Notably, when comparing NLSF and RSF solutions in LiBr (fully denatured) at similar concentrations a drop in this magnitude is observed (fig. S19).

Thus, we hypothesised that coupled with the reduction in MW, RSF is also losing its NTD, which as noted previously, is highly implicated in the improved rheological properties of NLSF and NSF. To prove this hypothesis, we conducted proteomics analysis on the peptide fragments (0.1-6 kDa) obtained directly during the dialysis of the liquid solutions of both RSF and NLSF. It was noteworthy that although substantial material was obtained from RSF, no material was recovered from NLSF. Together, the SDS-PAGE results obtained from NLSF, and this observation would indicate a near-native MW and the likely presence of all its functional domains. On the other hand, LC-MS/MS obtained from the RSF fragments found a total of 113 peptides matched the provided sequences (FibH, FibL and P25). With 35 matching FibH for 10.17 % coverage, 52 peptides FibL for a 96.56 % coverage, and 26 matched P25 with total coverage of 89.09 % (fig. S20, A, B and C). For brevity, the following discussion will be focused on FibH, given the predominant role of this protein in the system. However, the high coverage of both FibL and P25 indicates a high degree of hydrolysis in RSF (*64*). Of the 35 detected peptides from FibH, 22 belong to the first 150 amino acids; 2 of these have a 100 % confidence match, with other 3 high confidence matches with 81, 78, and 60 % of confidence. The higher number of matched fragments and their higher confidence alludes to preferential cleavages at the NTD (fig. S20, D and E). Although the results do not allow us to unambiguously determine NTD as the sole determinant for the transitions, as P25 and FibL could play a role in assembly, the high degree of NTD homology and presence across all *Ditrysia* (*51, 65, 66*), and the absence of the latter two proteins in Saturniids (*15, 67, 68*), reinforces the idea that this domain is critical in driving assembly.

Beyond this observation, the complete sequence coverage plot indicates that most of the matched peptides correspond to the more hydrophilic segments of the chain (fig. S21 A), i.e. hydrolysis during degumming in Na_2_CO_3_ is likely limited by accessibility. Furthermore, besides G, the most abundant residues in FibH, S and T are particularly enriched at the terminal positions suggesting possible intramolecular nucleophilic attack under the alkaline conditions. Other amino acids such as N, K, and E were found at the terminal position of the peptides. Curiously N, despite its low abundance with only 20 in FibH, was found predominantly at the C-terminal of the fragments, with S and T more abundant at the N-terminal positions (fig. S21 B). Overall, during degumming chain scission occurs mainly at S residues, as suggested by other authors (*69*), but other hydrolysis mechanisms are also occurring. The hydrophilic spacers and termini are richer in polar amino acids, often considered better nucleophiles in aqueous conditions. Residues R, H, and K are only found in the terminal domains, and E and D both in the spacers and terminals (fig. S21).

### Structure and supramolecular ordering

Despite the significant rheological differences observed with pH changes for NLSF, we did not observe clear differences in the NMR and CD data in the studied pH range, suggesting that most of the secondary structures of the protein remain unchanged. To better understand the system, we then conducted TEM of samples at pH 8 and 6, above and below the identified transition. Here we observed that the protein appeared as globules with an apparent size of about 24 ±8 nm at pH 8, but elongated fibrillar structures were observed at pH 6 (fig. S22 A and B, respectively). Although these fibrillar structures were slightly thicker than the expected solenoid (6 ± 1 nm against ca. 3 nm), acquiring higher resolution images proved difficult given their small size and high sensitivity of the samples to beam damage. The observation of no change in NMR and CD data with pH, but such dramatic morphological change observed in TEM suggests that only small subdomains withing FibH might be changing (such as NTD and the flexible linkers). This would allow for extension of the solenoid, which seems to be contained within the globules, suiting the observation in DLS and offering a conciliating explanation between the micelle and liquid crystalline models (*70, 71*). Beyond the morphological change, we observed the presence of supramolecular structures resembling bottle-brush structures (fig. S23) similar to those observed previously from *Samia ricini* (*Saturniid*) fibroin (*72*), and recently in NSF (*73*). Based on these observations and the apparent role NTD has on the reversible sol-gel transition, we propose that FibH can form high-order oligomers driven by head (NTD) interactions, making the core of the supramolecular fibre.

The structure of the NTD solved by X-ray crystallography has a saddle shape, with highly hydrophobic patches on the surfaces, proposed to form tetrameric units (dimer of dimers). The remaining β-sheet surfaces are covered by a short α-helixes linked to the rest of the structure by a flexible spacer (Fig. 4. D). Given that this helix can flip out of the hydrophobic space (*51*), we propose that NTD tetrameric units could stack leaving the rest of the multidomain protein body protruding laterally. These structures impart great ordering by creating supramolecular brush-like fibres (Fig. 4. E, F) which might be responsible for the cholesteric textures observed previously (*29*). Hence, we conducted docking simulations from NTD crystal units at pH 8, 7 and 6 to verify that these can undergo pH-dependent oligomerisation (fig. S24 A). Our results suggest that NTD likely forms higher-order oligomers, with the tetrameric interactions being strongly favoured at pH below 7 (fig. S24, B-E). At this pH, the protonation of residues Asp44 and Asp89 promote the stability of the interaction; both strictly conserved throughout all FibH in *Ditrysia* and belonging to acidic clusters (*66, 74*). Moreover, it was noted that all the interfaces (dimer/dimer and tetramer/tetramer), even if slightly different in estimated ΔG values, were in a similar order of magnitude. Current work is undergoing to obtain high resolution experimental data of this transition. Similarly, we noted that solenoid structures would be left in position for lateral interactions to emerge upon oligomerisation.

**Fig. 4.**
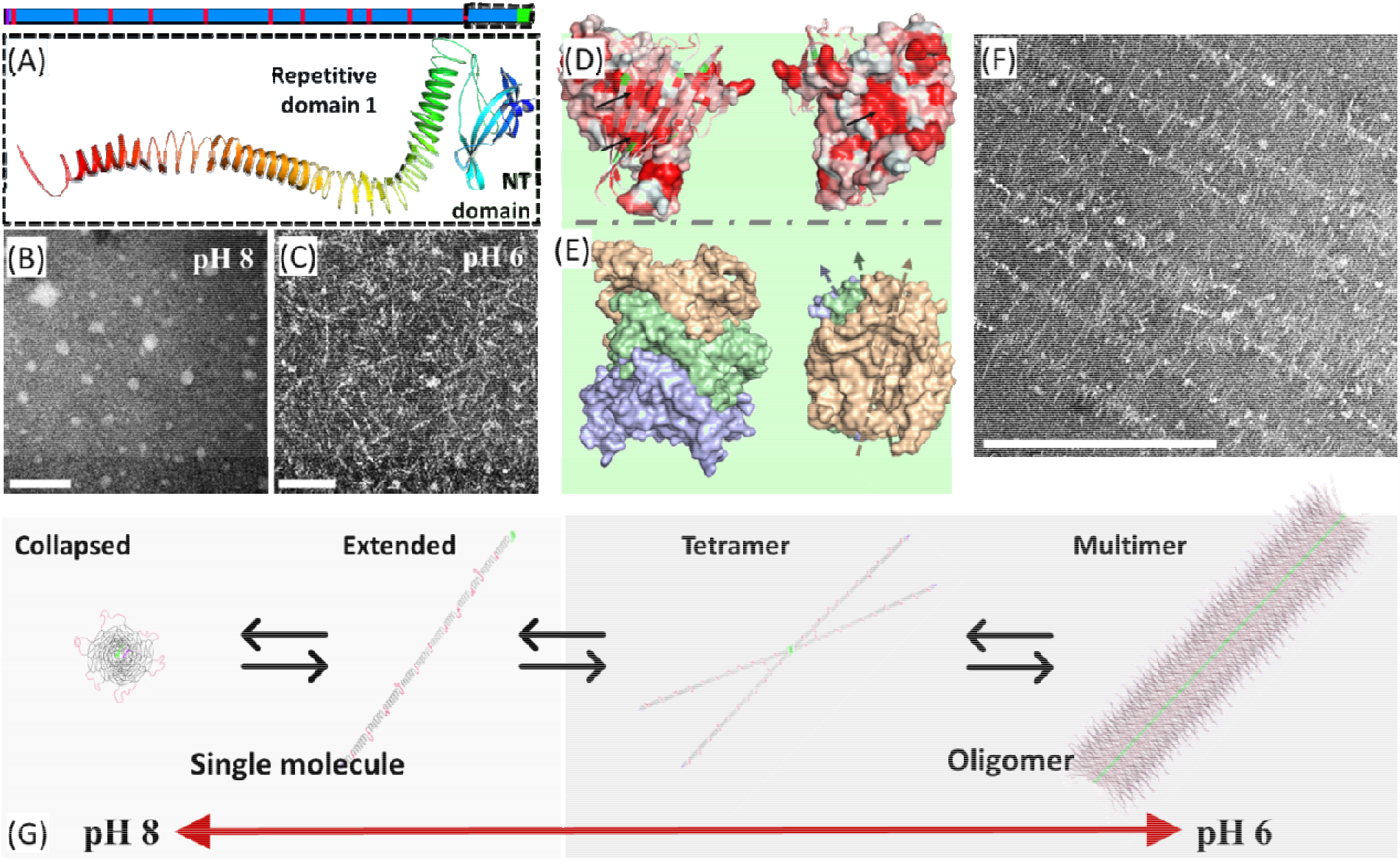
Evidence of a reversible pH-driven assembly. (A) Cartoon of Fibroin heavy Chain with 12 repetitive blocks (blue), 11 flexible linkers (red), NTD (green) and CTD (purple), under this a molecular cartoon of one of the folded models corresponding to NTD and first repetitive domain in a beta-solenoid type of fold. (B) Negative stained TEM of NLSF at pH 8. (C) Negative stained TEM of NLSF at pH 6. (D) Surface representation of the proposed biological unit of the NTD tetramer coloured by hydrophobicity with top dimer represented as a cartoon (left) and as seen from the back (right) with black arrows pointing at hydrophobic patches. (E) Proposed NTD tetramer stacking shows 3 tetrameric units stacked from the side (left) and the top (right) with arrows indication the twist of the stacking. (F) Negative stained TEM of NLSF at pH 6 showing supramolecular brush-like structures formed via NTD stacking. (G) Proposed pH driven self-assembly of the protein, from globular unimer, extended solenoid, dimer/tetramers and multimeric brush-like fibril (left to right). Scale bars 100 nm for B and C, and 500 nm for F.

Docking simulations of solenoid units verified that such interactions are possible and equally favourable parallelly or antiparallelly driven by the exposed A residues on the surface (fig. S25), providing a foci for network formation (fig. S26).

These structures likely fulfil an early role in the assembly pathway, given the conservation of the NTD domain across all species. However, no evidence of these bottle-brush structures can be found within the fibre (figs. S27 and S28) suggesting that these may exist transiently. NTD-driven oligomerisation facilitates controlled phase separation, thus aiding in the dehydration of the dope and fostering lateral interaction of the solenoid units. Upon acidification, these structures align with the flow generating the cholesteric textures. Here, extensive lateral interaction forms a network. Thus, in these first steps, the system is characterised by two main interactions, NTD-NTD and solenoid-solenoid; the absence of CTD and FibL in *Saturniids* indicates that these are not essential for fibre formation, as has been suggested recently (*75*). As the silk duct diameter is reduced, a critical stress is reached, and NTD interactions are disrupted, leaving a network dominated entirely by lateral solenoid interactions (fig. S29). The order imparted by the NTD oligomers is lost, and so the network suffers a reorganisation, with the solenoid axis aligning with the flow and giving rise to the recently observed fractal network, both in NSF and high-quality RSF (*70*). Fig. 5 summarises the proposed fibre formation model, with illustrations and experimental microscopy data showing snapshots of the observed structures and correlation with the optical textures.

**Fig. 5.**
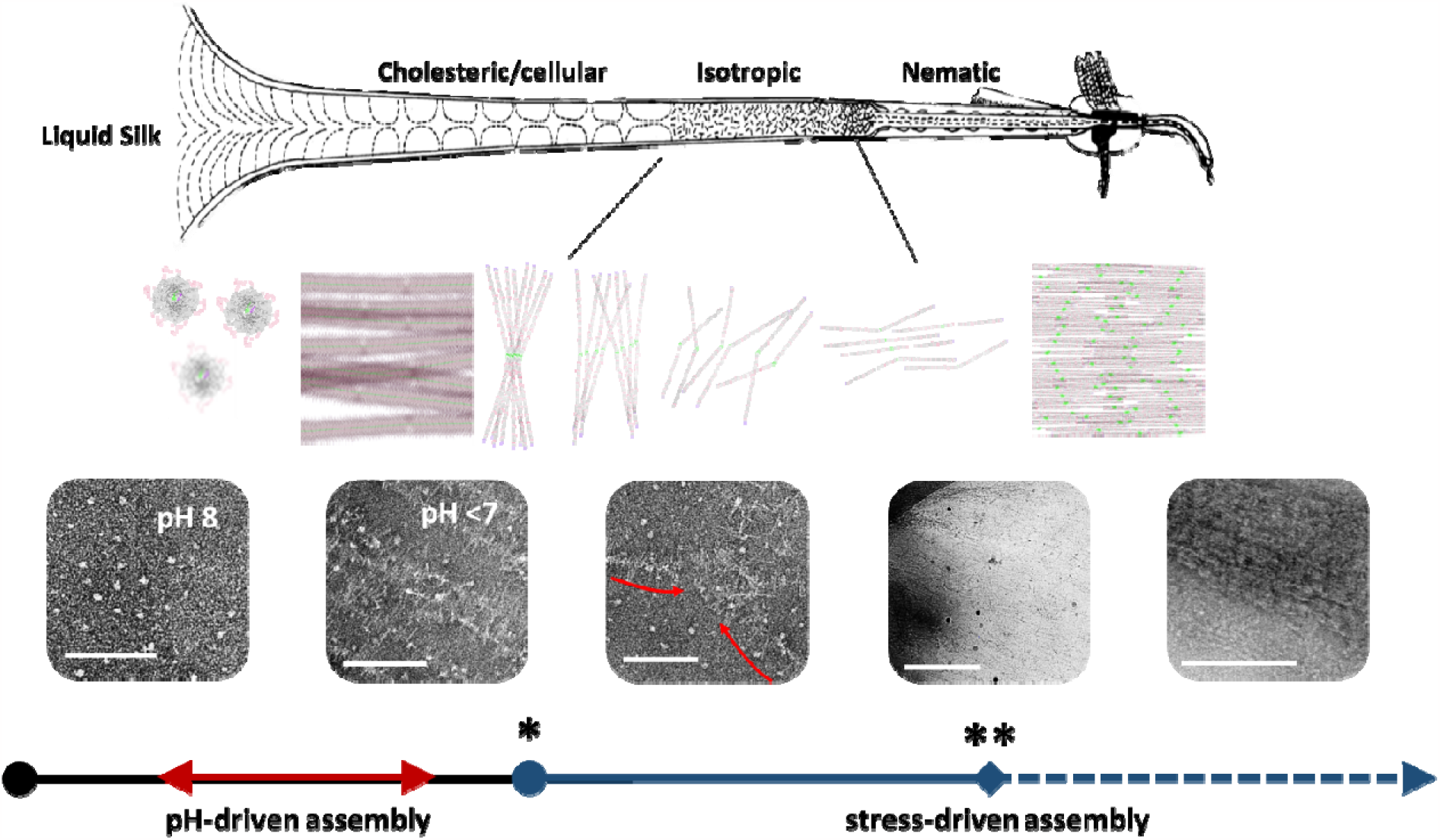
Summary of the newly proposed assembly pathway for fibroin into the silk fibre. Cartoon with the schematic representation of the proposed self-assembly pathway for fibroin. Illustration with optical textures was adapted from the literature.(*22*) In the first instance, fibroin at pH greater than 7 exists as a globular unit. At pH lower than 7 extend and form supramolecular brush-like fibrillar structures that would align with the direction of the flow and originate a cholesteric texture. At critical stress, the supramolecular structures break, leaving free isotopically ordered anisotropic oligomers of lower order that align and deform under flow to generate the solid silk fibre after the silk press.

The described transition explains the transformation of cholesteric-isotropic-nematic observed before (*27*). Moreover, the last step leaves the solenoid axis, and thus the hydrogen bonding network, parallel with the direction of elongation and ideally placed for unfolding and stretching of the backbone. Similar denaturation of α-helix fold of fibroin in 1,1,1,3,3,3-hexafluoro-2-propanol (HFIP) under stretching has been observed recently, with a α-to-β transformation prompted by stretching (*76*). These hypotheses are supported by observing two maxima in yield stress in stress-controlled experiments on gel-like material at pH 7 (fig. S30). Similar stress maxima were recently observed when pulling fibres from other native-like fibroin solutions (*77*). The first yield occurs at relatively low strains and appears narrower than the second. Such a double yield behaviour suggests the disruption of two different interactions, which we believe correspond to the initial disruption of NTD interactions, with narrower energy distribution, and secondly, a stress-induced denaturation (hydrogen bond breakage) with subsequent backbone stretching, similar to the critical amount of work observed for fibrillation (*77*). The described process leads to the extension of the backbone and β-sheet formation with “nano-fishnet” architecture proposed before (*79*). Based on this hypothesis, we would predict that if the stress is kept low enough during fibre formation, one would potentially be able to obtain Silk-I based fibres (or a combination of polymorphs), in agreement with the observation that the rate of work, and not the amount of work in itself are essential to promote the transition to solidification (*80*).

## Conclusions

This study employed a comprehensive array of techniques to enhance our understanding of the Silk-I structural polymorph of silk fibroin. Through our investigations, we made significant advances in understanding the in-solution structure of FibH and uncovered a pH-induced reversible sol-gel transition driven by NTD interactions, which we subsequently correlated with morphological and structural alterations of the protein at the molecular and supramolecular level.

We propose that FibH from *B. mori* is a multidomain protein with twelve low complexity domains with the ability to fold into β-solenoids. We accredit the Silk-I diffraction pattern to these domains, which are linked by flexible spacers and flanked by two terminal domains, of which NTD is the main driver of supramolecular assembly. As the pH drops below 7, NTD oligomerises forming large bottle-brush supramolecular fibres. The solenoid units, anchored within the construct are enabled to form stable lateral interactions driven by hydrophobic Ala-rich surfaces. Stress in the first instance disrupts the initial bottle-brush structures, leaving behind a fractal network of solenoids interacting laterally. As stress increases, the fold is denatured and the backbone of the polypeptide is stretched in the direction of the flow, driving the assembly into a nano-fishnet molecular architecture with β-sheet crystallites as nodes (*79, 81*). On the other hand, our study on standard RSF showed that these have largely lost NTD, FibL and P25. Thus, it contains mainly a polydisperse mixture of the low complexity domains of the protein. In the absence of NTD, these fragments are devoid of the pH switch, and therefore lack the degree of preorganisation, albeit still retaining most of the Silk-I features. Our mass spectrometry analysis of proteolysis can therefore provide a powerful method to assess feedstock quality.

Although the β-solenoid structure might be common among silk proteins, we believe it to be incidental, as other fibrillar structures are compatible with the proposed assembly mechanism. NTD being the only essential feature in pH-driven assembly. In silk-spinning *Samia ricini*, the repetitive domains are rich in polyA motifs folding in α-helixes (*82, 83*), and also form the bottle-brush structures (*72*). Therefore, our model provides a general description for silk-fibre formation independent of specific sequences. Beyond insect silk, our work indicates that other low complexity proteins, especially those rich in GX motifs, such as the class III of G-rich proteins and other non-characterised fibroin heavy chain-like proteins widely found across all kingdoms of life. We also note the uncanny resemblance of FibH with known toxic GA repeat proteins associated with disease (e.g. amyotrophic lateral sclerosis, ALS), which in the monomeric species might have a similar fold, and their aggregation pathway might be similar as the proposed here.

In a wider context, our work opens the opportunity for further interrogating the structure of simple repeat proteins, associated both with health and in disease development. Many of these are believed to be disordered, but superstructures of polyglycine-II or polyproline-II helixes might underpin their function. Within the silk community, our work also paves the way for the generation of a new class of materials based on native-like silk fibroins (NLSF). Overall, through our findings, we provide a unifying model that accounts for the observations on silk-fibre formation made through decades of research on the fibroin assembly process, from a liquid state to the remarkable structural material that constitutes silk fibre and might serve as inspiration for the design of novel bioinspired materials.

## Supporting information

Electronic supplemental information

## Acknowledgements

The authors would like to express their gratitude to the following individuals and facilities for their contributions and assistance: J.C. Eloi from the Chemistry Imaging Facility, J. Mantell from the Wolfson Bioimaging Facility, U. Borucu from the GW4 Cryo-EM Facility for their help and insightful contribution in TEM; M. Crump, and specially C. Williams for their assistance and discussions on protein NMR, and access to the 700MHz spectrometer; C. Arthur for contributions and assistance with LC-MS/MS; R. Cruz-Samperio for help with SDS-PAGE and discussions; P. Laity for contributions and thoughtful discussions related to this manuscript.

## Funding

EPSRC National Productivity Investment Fund grant EP/R51245/XF (R.O.M.T.).

EPSRC Doctoral Prize Fellowship at the University of Bristol grant EP/W524414/1 (R.O.M.T.). Wellcome Trust grants 086906/Z/08/Z, and 100917/Z/13/Z (N.S. and R.W.).

The EIC Accelerator grant 947454 (N.S. and R.W.).

The NIHR i4i Invention for Innovation award II-LB-0417-20005 (N.S. and R.W.). ^†^

EPSRC early career fellowship grant EP/S017542/1 (F.P.).

EPSRC, grants EP/K035746/1 and EP/M028216/1 (TEM).

Wellcome Trust grants 202904/Z/16/Z and 206181/Z/17/Z (TEM).

BBSRC grant BB/R000484/1 (TEM).

BrisSynBio, a BBSRC/EPSRC Synthetic Biology Research Centre, grant BB/L01386X/1 (NMR).

BBSRC Alert 20, grant BB/V019163/1(NMR). China Scholarship Council (Y.L.).

^†^The views expressed in this work are those of the author(s) and do not necessarily reflect those of the NIHR, the Department of Health and Social Care or any of their funding bodies.

## Author Contributions

Conceptualization: ROMT, SAD, LS, CH, NS, RW

Methodology: ROMT, SAD, CH, LS

Investigation: ROMT, LS, YL

Visualization: ROMT, YL

Funding acquisition: SAD, NS, RW, ROMT

Project administration: ROMT, SAD

Supervision: SAD, NS, RW

Resources: NS, RW, YL, FP, ROMT

Writing – original draft: ROMT, SAD

Writing – review & editing: ROMT, YL, FP, NS, RW, LS, CH, SAD

## Competing interests

The authors declare no competing interests.

## Data and materials availability

All data are available in the main text or the supplementary materials.

## Notes

### Competing Interest Statement

The authors have declared no competing interest.

### Summary of Updates

Updated text version. Simplified ELectronic supplementary information

