## Supplementary material for "Silk Road Revealed: Mechanism of silk fibre formation in *Bombyx mori*": Electronic supplemental information

Materials and Methods

Native-like silk fibroin solution (NL-SF)

Fibroin solutions of 8.5 M LiBr and 20 wt% of protein (proprietary protocol) were kindly provided by Orthox LTD in 60 mL syringes and stored at 4 °C until used. A stock solution at 2 wt% was made for studies requiring diluted solutions by diluting the provided solution with 8.3 M LiBr. The 20 wt% solutions look clear, free of precipitates, with a distinctive pale-yellow tint and low-shear viscosity of 60 ± 3 Pa.s. The stock solutions were then dialysed against MilliQ water using 12-14 kDa molecular weight cut-off (MWCO) dialysis membranes, with regular medium- to fresh MilliQ water changes. LiBr removal was followed by conductivity measurements of the dialysate and contrasted against a conductivity calibration curve, often requiring 3-4 days of dialysis to remove 99.9% of LiBr salt. After dialysis of the 20 wt% solution, a clear gel is obtained. In contrast, the 2 wt% yields a clear liquid solution with viscosity like pure water. Concentrations were determined gravimetrically by cutting out small fragments or aliquoting about 500 µL of the solution and lyophilising them overnight in a Labconco Freezone 2.5 L Freeze-drier. The high concentration solution produces a gel with concentrations in the range of 60-80 mg/mL, whereas the 2 wt% solution produces a solution in the range of 5-7 mg/mL.

Standard Regenerated silk fibroin solution (RSF)

A standard regeneration protocol was followed (*30*). Briefly, about 10 g of raw silk, kindly provided by Orthox LTD, were boiled for 30 min in 4 L of aqueous 0.02 M Na_2_CO_3_, the supernatant was then discarded, and the fibres were rinsed with copious MilliiQ water at room temperature. The wet degummed fibres were dried in a convection oven at 30 °C overnight. The dry degummed fibres were weighted to verify degumming efficiency. Finally, degummed fibres were dissolved by adding specified volumes of 9.3 M LiBr to obtain a 20 wt% silk fibroin solution and left in an oven at 60 °C for 4 hours, or until total dissolution was observed (whichever occurred first). The solution was then centrifuged at 5000 units of relative centrifugal force (RCF) for 5 minutes at room temperature to remove small undissolved particles. After these steps, the solution looks clear, free of precipitates with similar pale-yellow tint and with low-shear viscosity of 2 ± 0.5 Pa.s. The solutions were then dialysed against MilliQ water following an identical procedure as before. After dialysis, the 20 wt% solution produces a clear solution with a viscosity not different to pure water. Concentration was determined gravimetrically by aliquoting 500 µL of solution and lyophilising it overnight. The 20 wt% LiBr solution produces an aqueous solution with 60-80 mg/mL. Given the liquid nature of this sample, dilutions were prepared directly from the concentrated stock.

Sodium Dodecyl sulphate Polyacrylamide Gel Electrophoresis (SDS-PAGE)

For SDS-PAGE measurements, Tris-acetate 8-12% gels were used from Sigma. Samples of 0.1 mg/mL of NL-SF and RSF were used by mixing with stock solutions of running buffer and coomassie blue. A high molecular weight ladder was used obtained from , with reference proteins: .Gels were run as recommended, using 200 V and 1 Amp for 40 minutes. Gel images were then analysed using FIJI by measuring grey-scale values and plotting coaligned with the standard ladder.

Rheology

The rheological characterisation was done in a Malvern Kinexus Pro rheometer with a Peltier lower plate for temperature control and with either a 20 mm parallel plate (PP20) upper geometry for gel samples, or with a truncated conical geometry with 4 degrees angle and 40 mm in diameter (CP4/40) for liquid samples. Samples were equilibrated for 24 hours at room temperature in x50-100 volume of 100 mM Tris buffer at indicated pH values, titrated using stock solutions of 2 M HCl or 5 M NaOH. Although the buffering efficacy of Tris at pH 6 is very low, this buffer is preferred as other common buffers such as phosphates reduces the storage time considerably, promoting early conformational changes to the NL-SF concentrated solutions/gels.

Briefly, 500 mL of buffer solution were prepared at the desired pH. About 5-10 mL of aqueous SF solution were placed in them, contained by similar dialysis tubing membrane from previous steps. Samples were then left under slow stirring (ca. 150 rpm) at room temperature for 24 h to equilibrate. Following this, samples were taken out from the buffer solution and either carefully poured into clean 50 mL conical end centrifuge falcon tubes for liquid samples or taken to standard plastic Petri dishes (100 mm diameter) for gel samples. These last samples were taken out of the dialysis tube by carefully cutting open the membrane with a clean new scalpel. Samples were used immediately afterwards.

The following procedures were adapted from different works in the literature that work with native silk fibroin solutions, with minor modifications. In every case, samples were treated with the utmost care to prevent early conformational changes prompted by poor handling. Gel phase samples were carefully collected by excising small fragments from the bulk using a wet scalpel and disposing of any sample that would show evidence of aggregation (opaque fibrillar aggregates). Enough gel sample was used to fill the entire geometry, carefully positioned in the centre of the lower plate, with the upper plate lowered slowly with a maximum allowed normal force of 1 N to a 1 mm gap. Water saturated tissue was left around the geometry, making sure it was not in contact with it, and the entire geometry was enclosed by a plastic cover to prevent dehydration during experiments. For liquid samples, a 5 mL pipette with tips with cut ends were used to prevent applying high shears when handling, with a standard quick loading setting from the software and similar procedures to prevent dehydration.

For the determination of the Linear Viscoelastic Limit (LVL), both elastic (G’) and viscous (G”) moduli were measured in amplitude sweeps done in oscillatory mode at a constant frequency of 1 Hz, in the strain range between 0.1 and 100%, with 10 points per decade. Measurements at each point were taken until convergence was detected by the Kinexus software. Liquid samples underwent a pre-homogenisation step, as proposed elsewhere (*49*, *84*), by conducting fully rotational low-shear rate (1 s^-1^) deformation for 100 s. At least 3 repeats were done for every sample.

After determination of LVL, G’ and G” were measured under frequency sweeps on fresh samples at constant strain of 0.01 (1 %, within LVL of all samples) between 10 and 0.1 Hz. Similarly, 10 points per decade were measured, with measurements being taken until data convergence was observed by the software. Liquid samples underwent homogenisation, as mentioned before, by applying fully rotational deformation at 1 s^-1^ for 100 s before conducting the experiments. At least five repeats were done for every sample.

Only liquid samples were used for viscometry. Viscosity against shear rate measurements was obtained by a shear rate table in the range between 0.1 and 100 s^-1^, with 10 points per decade. Measurements were taken until convergence was obtained at each shear rate point. Shear viscosity, as well as normal force values, were recorded.

Fibre X-ray diffraction

These experiments were done using Rigaku rotating anode (Cu Kα) with Saturn CCD detector with exposure times of 30–60 s and specimen to detector distance of 50 or 100 mm. Reflections were measured using CLEARER software.

For diffraction obatained from degummed silk fibres, about 1 cm long fibre bundles were mounted on glass capillary tubes using epoxy-based glue and then mounted directly onto a goniometer head for alignment. Alignment was ensured by monitoring a microscope camera focused on the beam path, and alignment was adjusted for the nominal 0 and 90° orientations. Spectra were acquired at 50 and 100 mm of distance at 0 and 90° orientations, for 30 and 60 s, respectively.

The silk-I film was fabricated by slow drying about 100 µL of NLSF solution at 6 mg/mL at room temperature, drop-casted onto a clean petri dish surface. Once dried, the film was carefully removed using forceps, glued to a capillary glass tube, and observed as the fibres before. Here, the film was orientated so the beam would strike normal to the surface at 0°, parallel to the film surface at 90° and later a range of acquisitions were done by varying orientation by 5° between 0° and 90°.

Liquid Chromatography-Mass Spectrometry (LC-MS/MS) and Proteomics Analysis

Samples were analysed following modified Shotgun proteomics protocols. For the analysis of the hydrolysis patterns, concentrated solutions in LiBr of both samples were dialysed against MilliQ water while being contained within a dialysis tubing membrane with 6 kDa MWCO, which in turn was contained by a 100 Da dialysis tubing membrane. Care was taken not to cross-contaminate the different spaces, and the space between membranes was filled with 15 mL of MilliQ water. After 99.99 % of LiBr was removed by dialysis, determined by the conductivity of an 8 h dialysate from fresh, the samples collected from the volume between the 6 kDa and 100 Da membranes were transferred to sterile 50 mL conical centrifuge falcon tubes, from which samples were directly summited to be analysed. Obtained sample from RSF contained about 2.0 ± 0.5 mg/mL of peptides, but no material was recovered from NL-SF (< 0.1 mg/mL).

Samples were injected in a Dionex RS3000 High pressure Liquid Chromatographer (HPLC), coupled with a Orbitrap Elite in Electrospray ionisation mode (ESI) to analyse the fragmentation of the submitted peptides by ESI-MS/MS.

The collected data was then analysed using PeptideShaker and contrasted against the sequence of FibH, FibL, and P25 from the UniProt database (accession P05790, P21828, and P04148, respectively).

Nuclear Magnetic Resonance (NMR)

For NMR studies, only samples at pH 8 and 6 of NL-SF were studied. Briefly, stock 2 wt% solution was dialysed against MilliQ water until >99.9 % of LiBr was removed to obtain a solution of about 1 mg/mL of protein. These solutions were then passed through a NAP45 non-interacting column, following manufacturer instructions, pre-equilibrated with 10 mM Tris in deuterium oxide, titrated beforehand to the desired pH with aqueous 2 M HCl or 5 M NaOH. The collected buffered samples were then analysed using a Bruker Avance III HD 700 with a 1.7 mm cryo-enhanced probe. ^1^H-NMR were collected, as well as proton-proton Total Correlation Spectroscopy (^1^H-^1^H TOCSY) for residue assignment, Nuclear Overhauser Effect Spectroscopy (NOESY) for distance determination and natural abundancy carbon-proton Heteronuclear Single Quantum Coherence (^1^H-^13^C HSQC) for secondary structure determination of A and G residues.

Fold simulations and model generation

Fold simulations were carried over at two different servers to verify results congruency. However, in general, the AlphaFold2 prediction methodology was used. The services used were those published on Google Colab books by DeepMind and ColabFold teams (*34*, *35*). For the simulations, Colab Pro, was used to ensure higher assigned memories, better processors and longer runtimes.

*Fibroin Heavy Chain fold simulation*

Given the detected modularity of the multidomain protein and the size limitation on the simulations through ColabFold (1000-1400 residues), only the NTD and the first repetitive domain were used initially for a total of 650 amino acids. The multisequence alignment (MSA) was run using Jackhammer and mmSeqs2, with a minimum pair coverage with a query of 50% and minimum sequence identity of 20%. Jackhammer offered the best coverage overall, with 7456 Sequences found in Uniref90, 9062 Sequences found in SmallBFD, and 482 Sequences found in Mgnify, or 17000 sequences in total; most of these covered the repetitive domain. Next, Alphafold was run, and five models were generated after running the trunk of the AlphaFold neural network 8 times with different random choices across the MSA. Finally, the generated five models were relaxed using Ammber-relax forcefield, and the models were downloaded, visualized and post-processed using Pymol. On the other hand, mmSeqs2 found about 45 sequences, primarily for NTD, and the repetitive domain was modelled as a random coil.

Later, the same sequence was also run on the DeepMind colab book, and very similar models were obtained. Following these, the rest of the domains were modelled in fragments below 1000 residues that contained entire domains (either repetitive domains alone, in combination with one linker, or as two repetitive domains with a single linker). Models obtained through this method were used to generate the full FibH structure.

Docking simulations

*NTD docking*

In the first instance, the solved N-terminal tetramer model (PDB: 3UA0) was relaxed using Rosetta. Then, using the relaxed model, an octameric (tetramer-tetramer) stacked unit was generated with manually adjustments, and the model was further relaxed as an octamer.

Docking simulations under pH control were done by first determining the protonation state of charged residues on the relaxed octamer model. Next, protonation states were predicted by PDB2QR server (<https://server.poissonboltzmann.org/pdb2pqr>) at pH 6, 7 and 8. These generated models were used as references for root-mean-square deviation (RMSD) analysis during docking experiments. Finally, local docking was run in Rosetta on each of the three models, generating 2,000 independent simulations for each. Estimated free energy of interaction measured in Rosetta energy units (REU), ΔΔG, was then plotted against RMSD (a measure of the deviation from reference models).

*Solenoid lateral dockings:*

Models obtained from the first repetitive domain was used here. It was noted that the first repetitive domain could be divided into 3 sub domains linked by less regular solenoidal structures that are thought to allow for flexibility within the repetitive domain. Furthermore, the first and second subdomains were unique to the first repetitive domain, whereas the third subdomain was more represented across all repetitive domains. Hints at this were already observed after self-similarity analysis. Consequently, the model was separated into three sliced models containing the subdomains, each relaxed in Rosetta previous to global docking simulations, where only models with ΔΔG lower than -30 REU were used for further analysis.

Far UV Circular Dichroism (CD)

CD measurements were performed using low concentrations of protein (ca. 0.05 mg/mL) in 10 mM sodium phosphate buffer at the desired pH (6-9) at a temperature of 25 °C. Buffer pH value was adjusted with aqueous dilutions of phosphoric acid or NaOH. Samples from NL-SF were obtained by dialysing stock 2 wt% solution in 8.3 M LiBr and equilibrated in x100 volume of buffer over 24 h. After equilibration, samples were centrifuged at 6000 RCF for 10 minutes at room temperature to remove any precipitate and later diluted with fresh buffer until obtaining High-tension (HT) values under 700 for the whole studied range. Spectra were acquired from 240 to 190  nm in a 1-mm path length quartz cuvette using a JASCO J-820 spectropolarimeter. The wavelength step for the CD measurements was 1 nm, and the scan rate was 100  nm/min. The shown spectra were background corrected and averaged over five scans. For temperature analysis, similar procedures were followed, the sample was heated in 5°C steps from 25 to 95°C and equilibrated for 30 s at each temperature.

Dynamic Light Scattering (DLS)

DLS measurements were performed using 1 mg/mL protein solutions. After dialysing the 2 wt% in 8.3 M LiBr solution, samples are collected with approximate concentrations of 1 mg/mL, equilibrated in buffer at desired pH (10 mM sodium Phosphate) over 24 h and used directly after centrifugation at 6000 RCF for 10 mins. Disposable PS large volume cuvettes were used in a Malvern Instruments ZetaSizer nano ZS where data was acquired using software that included macros for size analysis and temperature trends. Software provided materials parameter values for protein sample, and water for media was used. For temperature trends, samples were equilibrated for 60 s at each temperature, 3 scans were averaged, and the temperature ranges from 25 to 75 °C with 1 °C increment.

Transmission Electron Microscopy (TEM)

Jeol JEM 1200 with a tungsten filament and acceleration voltage of 120 kV and a Jeol JEM 2100 with a LaB6 electron gun at 200 kV. Both were used for brightfield and negatively stained images.

Tecnai T20 with a LaB6 filament and acceleration voltage of 200 kV for negatively stained samples, and a Tecnai F20 with a field emission gun (FEG) with an acceleration voltage of 200 kV for cryo-TEM. For the latter, a Vitrobot Leica EM GP with humidity set to 100%, blotting time 1 s, was used after adding 5 μL of the sample before plunging into liquid ethane, and grids transferred to a cryo-holder (Gatan, Inc.).

The higher resolution cryo-TEM were taken using a Talos Arctica equipped with a 200 kV X-FEG, Ceta 16 M CCD detector, Gatan K2 DED, and Gatan GIF Quantum LS energy filter. Samples were vitrified in this case using a FEI Vitrobot.

Supplementary Text

NOESY

Within the entire sequence of FibH, Y is only found next to A a total of 13 times, out of the 275 Y residues present. However, the ^1^H- ^1^H NOESY experiments showed consistent NOE signals between the A βH and Y δ,εH (aromatic). It is known that NOE signal depends inversely on the interatomic distance to the sixth power; or alternatively, the distance is directly proportional to the intensity to the -1/6 power. Thus, generally for NOE signal to appear, the two atoms need to be within about 6 Å. In fact, after using the internal Y interatomic distances as mean of calibration, it was possible to estimate average distances between A βH and Y δH to be about 3.4 Å and between A and Y εH 3.3 Å. Although these might be underestimated. There are 221 Y residues within the repetitive domains in total, with an estimated of 74 motifs that would place Y next to A in an inter-strand motif as described in the main text (Fig. 2), thus after this adjustment, the estimated distance would be about 2.2-2.3 Å between the aromatic protons of Y and A βH, in an almost perfect agreement with the model (ca. 2.5 Å). Similar calculations can be made for the observed and expected NOEs from Y to S or V, where Y to S are expected to be similarly across strand distances at least 19 times, and Y to V intra-strand distances (motifs VGY) a total of 27 times, the expected distances are of about 2.4-2.8 Å and about 3.2-3.7 Å, while the corrected measured distances are 2.1 Å and 2.3 Å, respectively.

Polymer viscosity

For polymers beyond a critical entanglement concentration, the viscosity follows a power law proportionality to the molecular weight as shown in Equation 1.

$$\eta={kM}^{a}$$

Equation 1

Where $\eta$ is the shear viscosity, $k$ is a polymer dependent constant, $M$ is the molecular weight, and $a$ is the proportionality exponent, which for most polymers assumes the value of 3.5 with very low deviation for a myriad of polymers. So much so that the proportionality is often directly expressed with this value. Consequently, $\eta\propto M^{3.5}$ and thus, if one considers the relative molecular weight reduction observed in the SDS-PAGE, it is about half of the original MW. This reduction in MW would be translated in a proportional reduction of $\eta$ by $\left( 0.5 \right)^{3.5}$, or 0.088.

Furthermore, viscometry analysis of the samples, NLSF and RSF, was conducted in the LiBr solutions at 20 wt% of concentration. The standing assumption is that in the LiBr solution, the protein is in its statistical coil conformation as a polymer in solution. As it would be expected, under these conditions, the value of the zero shear viscosity ($\eta_{0}$) for NLSF and RSF are 58.1 ± 0.2 and 2.13 ± 0.02 Pa.s, respectively. One order of magnitude difference was predicted from the previous power law relationship. The fitting of the experimental curves was done using the Cross method (see equation contained within fig. S19), where viscosity as a function of shear rate ($\dot{\gamma}$), is fitted using the infinite shear viscosity ($\eta_{\infty}$), $\eta_{0}$, a relaxation time ($\tau$) and a power coefficient (*m*).
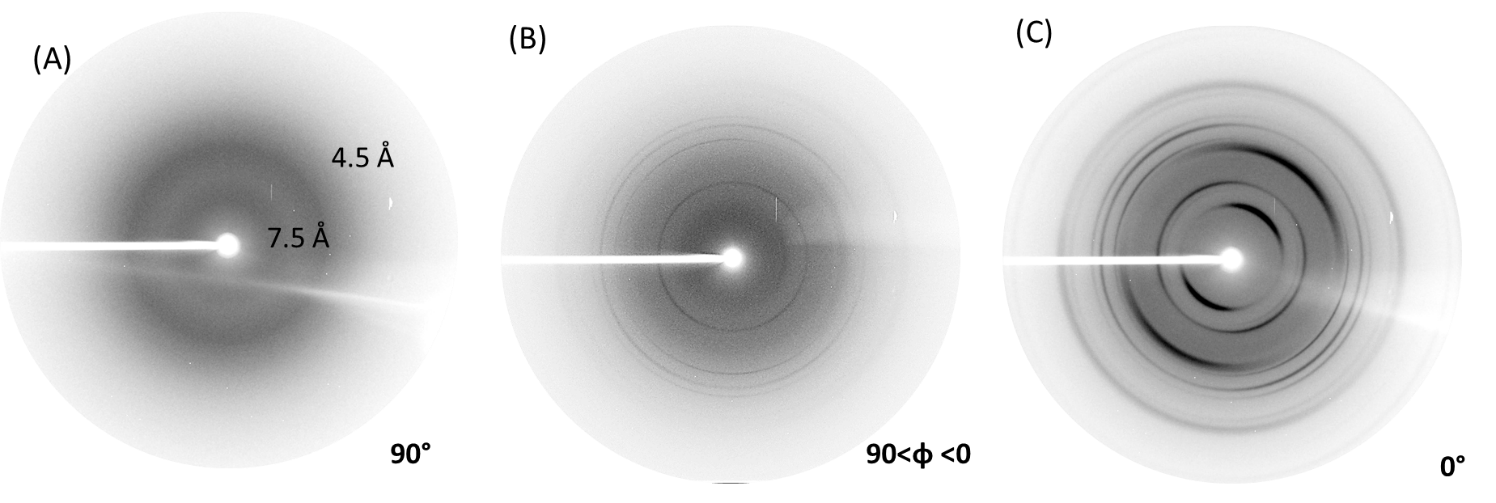

Fig. S1.

**2D X-ray diffraction in Fibre diffraction obtained from Silk-I film at different illumination angles relative to the film plane.** Diffraction pattern obtained at 0° (A), between 0 and 90° (B) and 90° illumination (B).

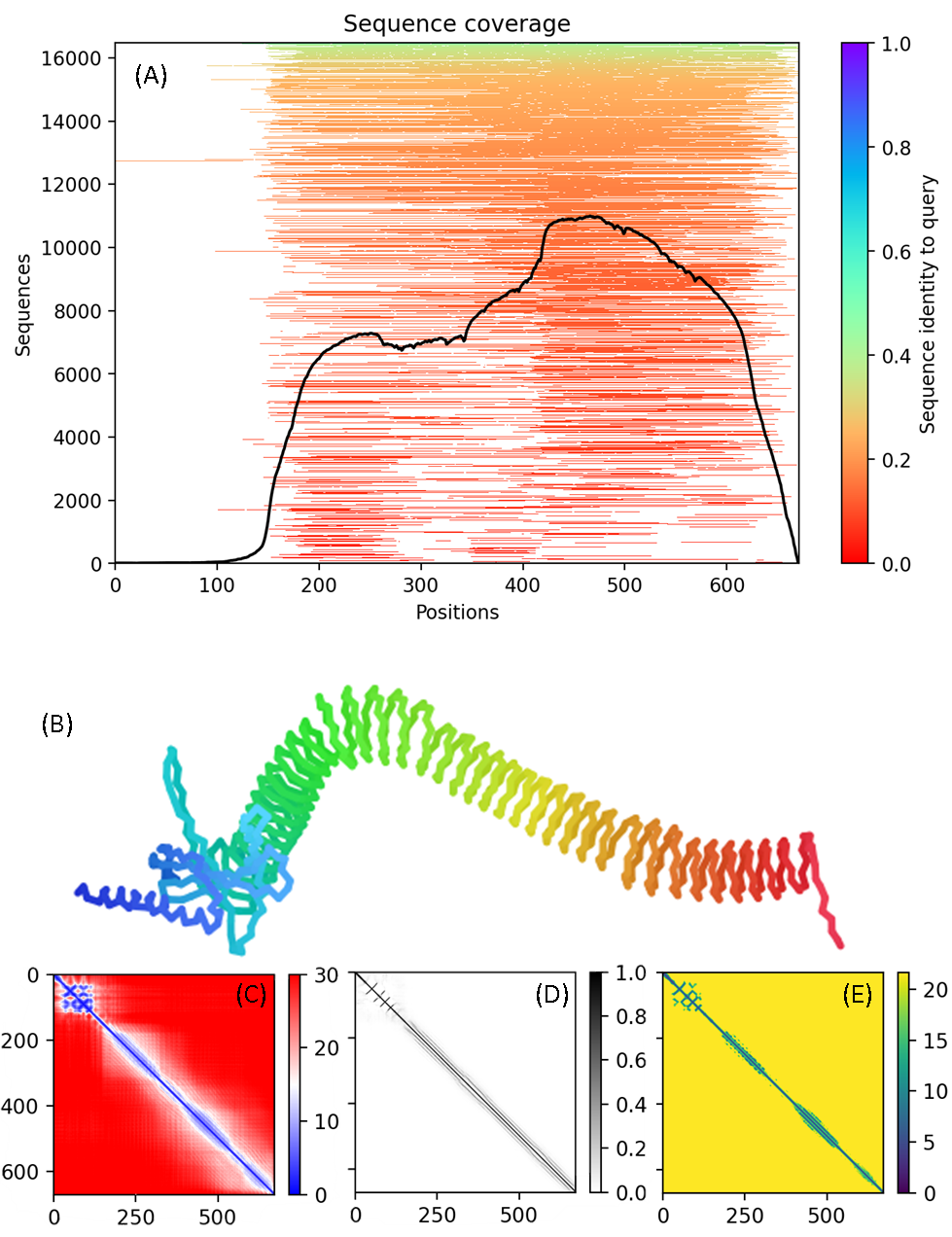

Fig. S2.

**AF2 simulation of NTD and first repetive domain.** (A) Multisequence alignment (MSA) results from a jackhammer database search showing +16000 found sequences aligned to the sequence of the first 650 residues of FibH (NTD+ first repetitive domain). (B) Ribbon representation of one of the five obtained models showing β-solenoid conformation coloured blue to red in an N to C direction. 2D plots of the predicted alignment error (PAE, C), predicted contacts (PC, D) and predicted distogram (PD, E) for the shown model. In the 2D plots, sequence residue position is plotted both on the horizontal and vertical axis, and the predicted value is in the out-of-plane axis, with the range and coloured coding being shown on the right-hand side of each plot.

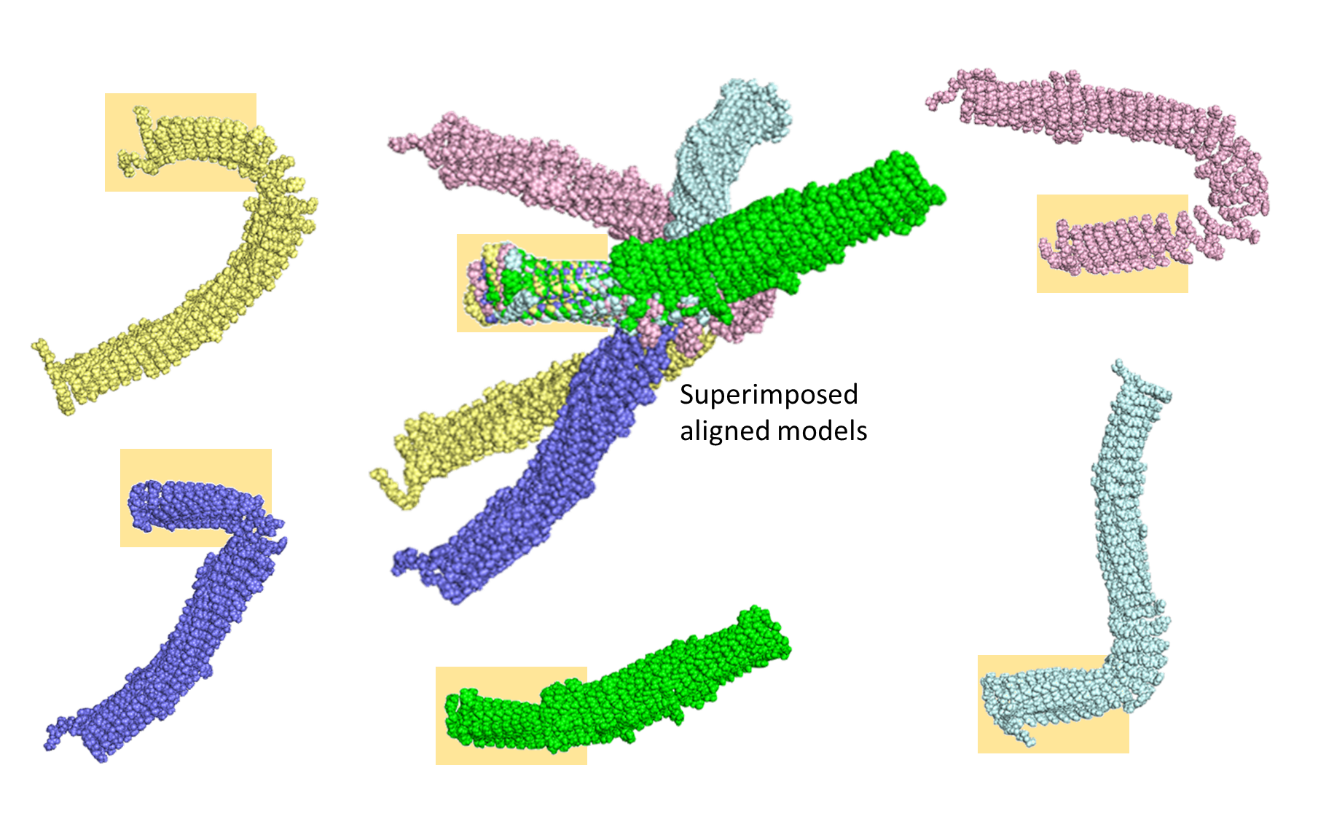

Fig. S3.

**Conservation of the β-solenoid topology for different generated models.** Schematic representation of the five obtained models for the first repetitive domain, aligned at the apparent first rigid sub-domain and superimposed (centre) to illustrate possible flexibility of the solenoidal structure.

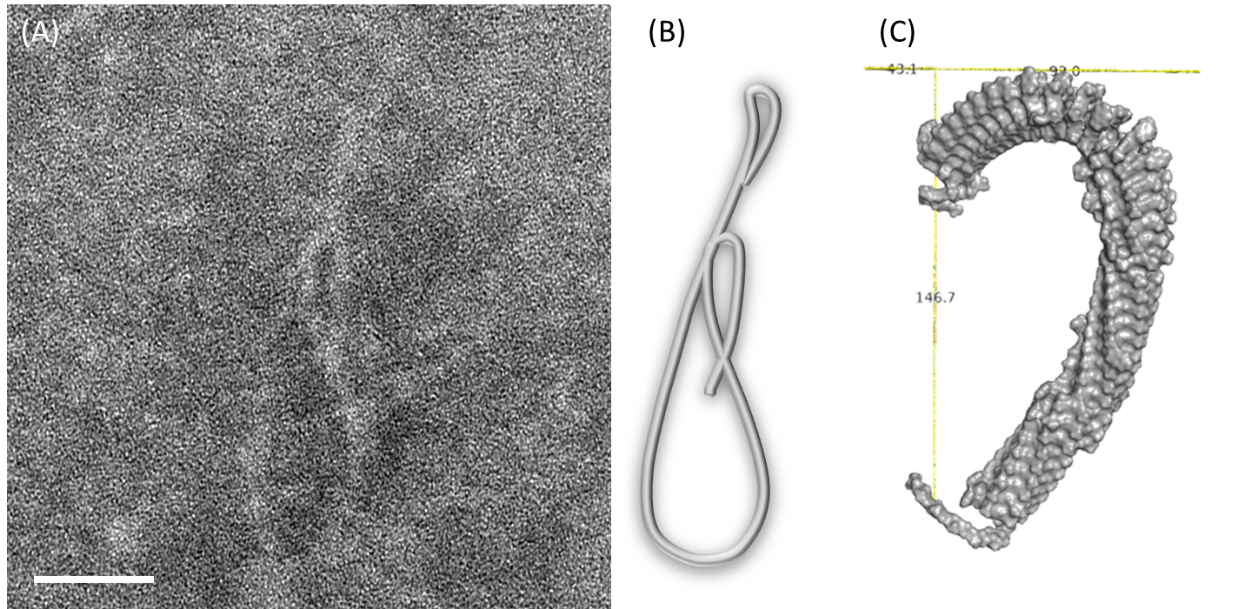

Fig. S4.

**Analysis and interpretation of TEM micrograph and FibH structure.** TEM image obtained from lightly stained NLSF sample at pH 6 (A), a drawing showing interpretation of the observed profile (B) and a surface representation of a segment of the obtained model from the first repetitive domain showing similar curvature to that observed in the image (C). Scale bar 40 nm.

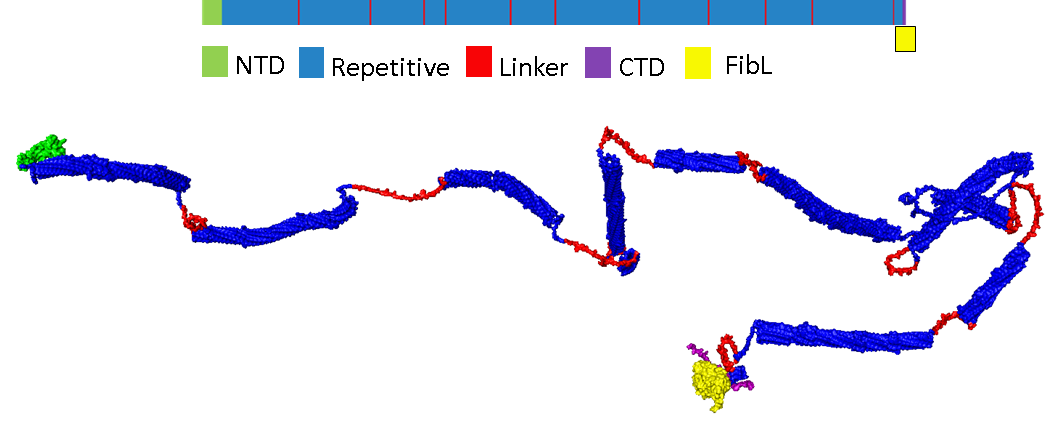

Fig. S5.

**Structure and topology of full FibH and FibL.** At the top, a simplified cartoon of FibH plus FibL showing the multidomain structure at the primary level, with a legend of domains and colouring shown below. At the bottom is the proposed secondary level structure with the domains coloured following the previous cartoon.

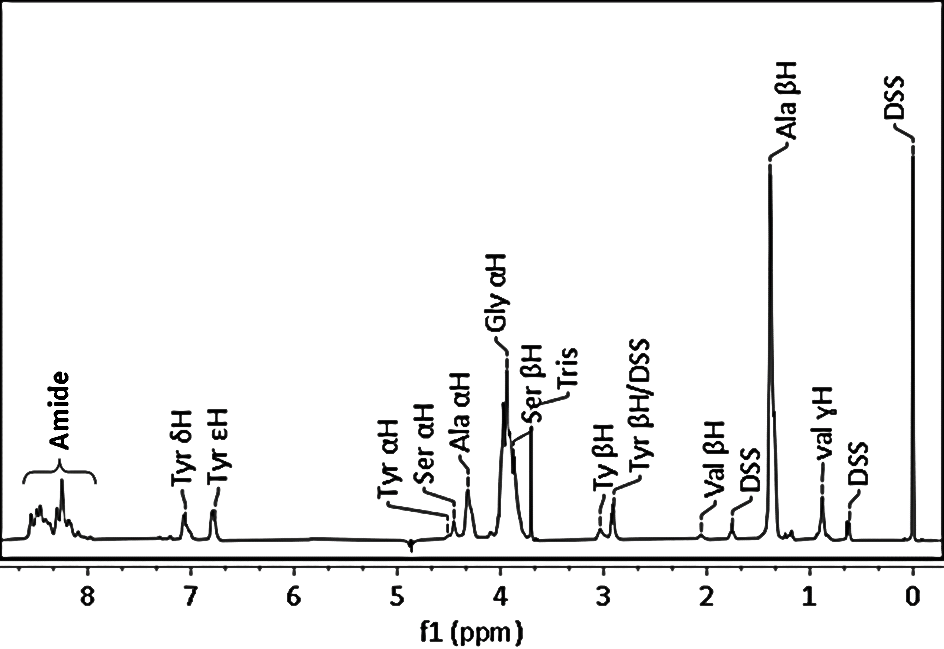

Fig. S6.

**One dimensional proton of fibroin**. Simple proton spectra of partially deuterated NLSF with chemical shift assignment ran using a Bruker Advance III 700 MHz instrument at pH 6 and referenced with DSS.

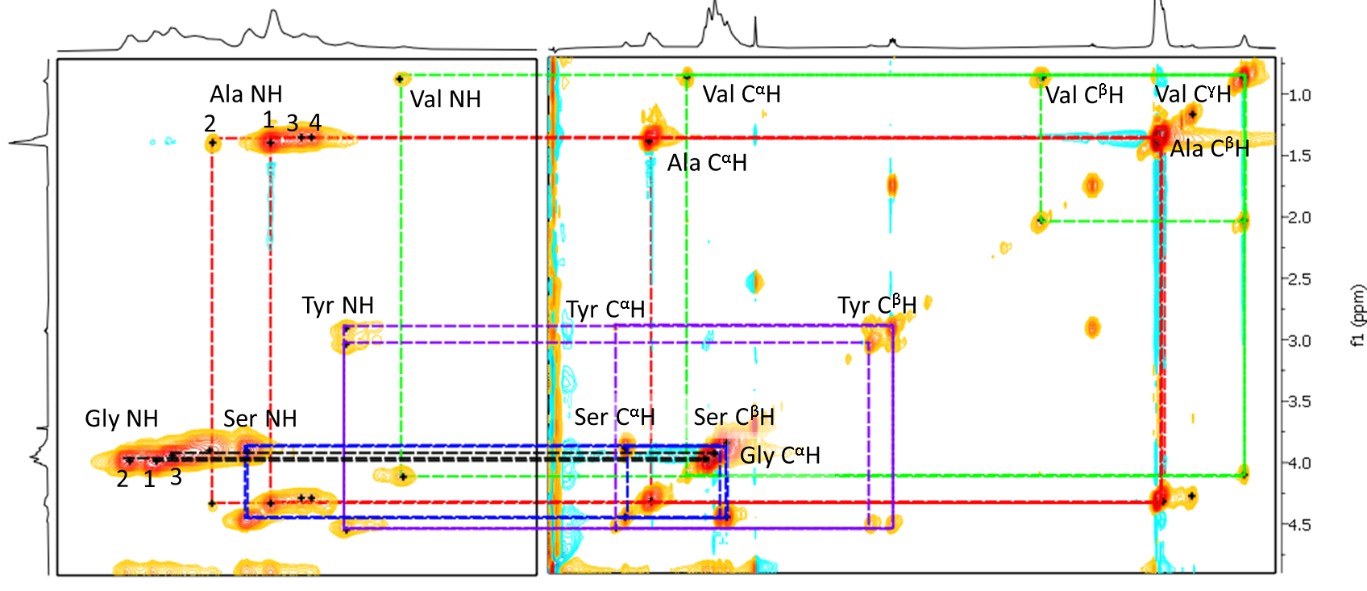

Fig. S7.

**Chemical shift assignment of proton**. Homonuclear ^1^H- ^1^H TOCSY experiment conducted using a Bruker Advance III 700 MHz instrument at pH 6 and referenced with DSS of NLSF (5 mg/mL). Chemical shift assignment is shown regarding the amino acid and coupling connections shown in coloured boxes/lines. Boxes/lines are coloured by amino acid, with G, A, S, Y, and V is shown with black, red, blue, purple, and green lines, respectively).

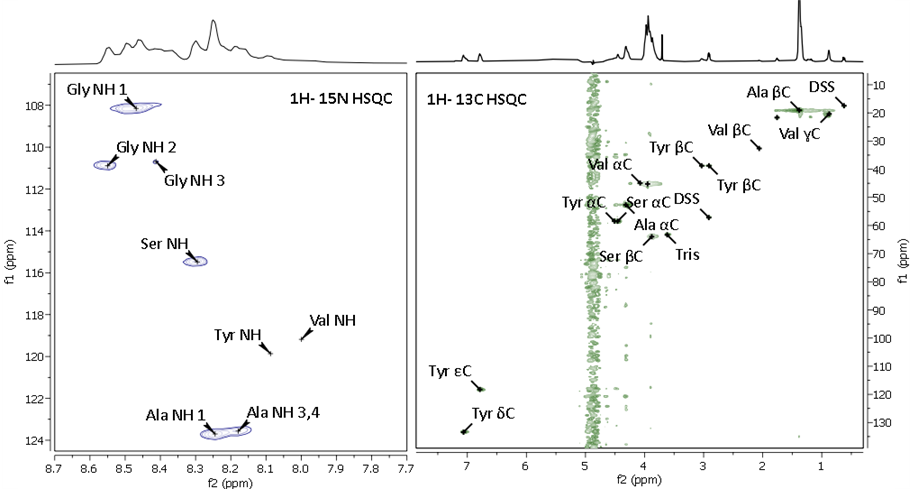

Fig. S8.

**NMR chemical shift residue assignment.** Heteronuclear ^1^H- ^15^N HSQC (left, blue) and ^1^H- ^13^C HSQC (right, green) with their respective shift assignment.
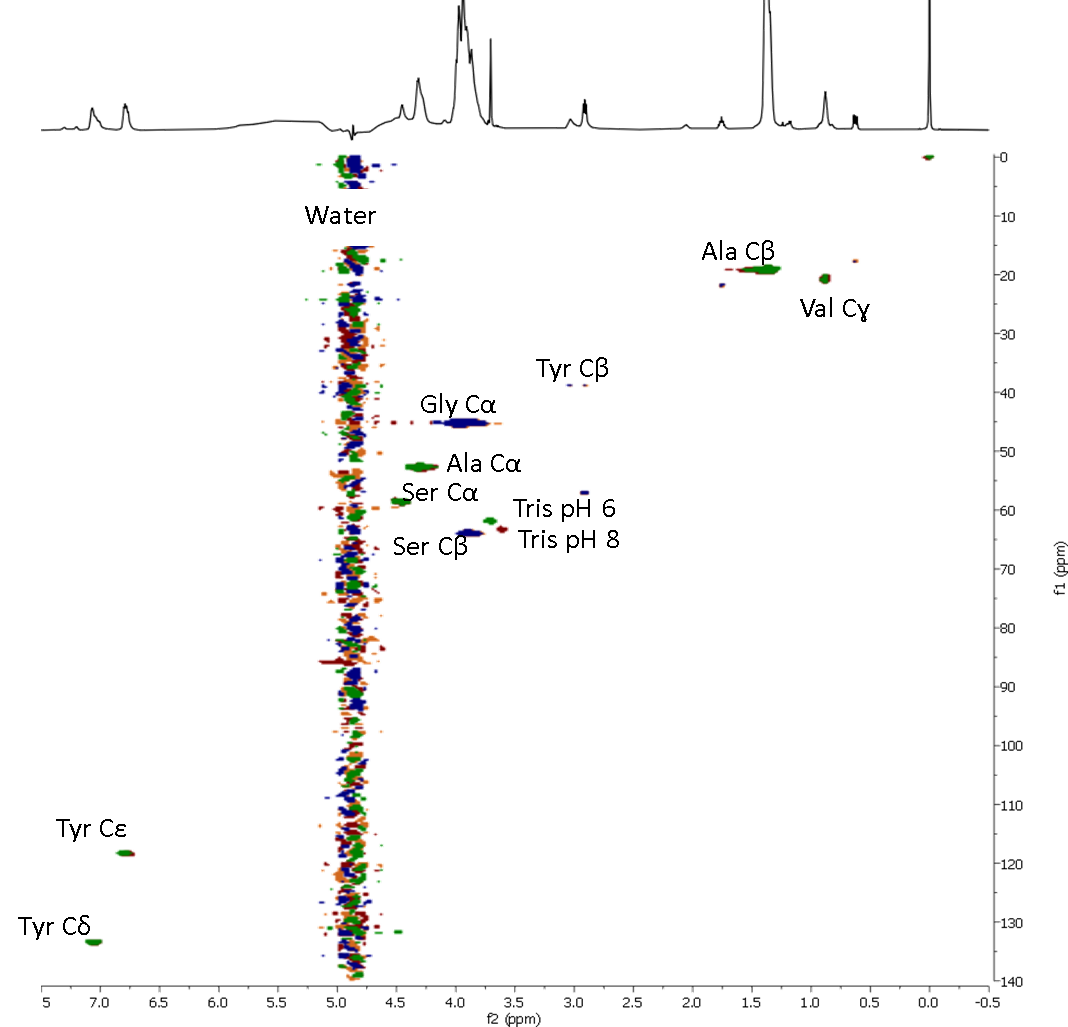

Fig. S9.

**Effect of pH on chemcial shifts.** Superimposed ^1^H- ^13^C HSQC spectra obtained for NLSF samples buffered at pH 6 and 8, showing no change in the chemical environment of the detectable residues.
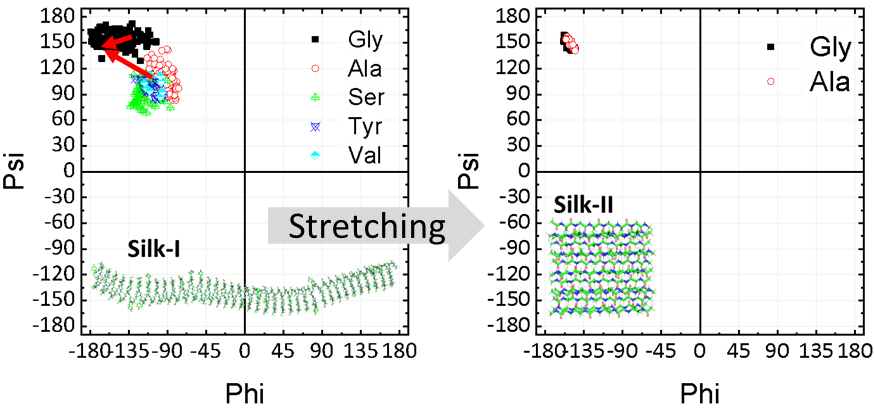

Fig. S10.

**Silk I to Silk II transformation, dihedral angles perspective.** Proposed stretching induced transformation of the proposed Silk-I to the theoretical Silk-II (DOI: [10.5452/ma-cs24y](https://dx.doi.org/10.5452/ma-cs24y)) as observed from dihedral angles.

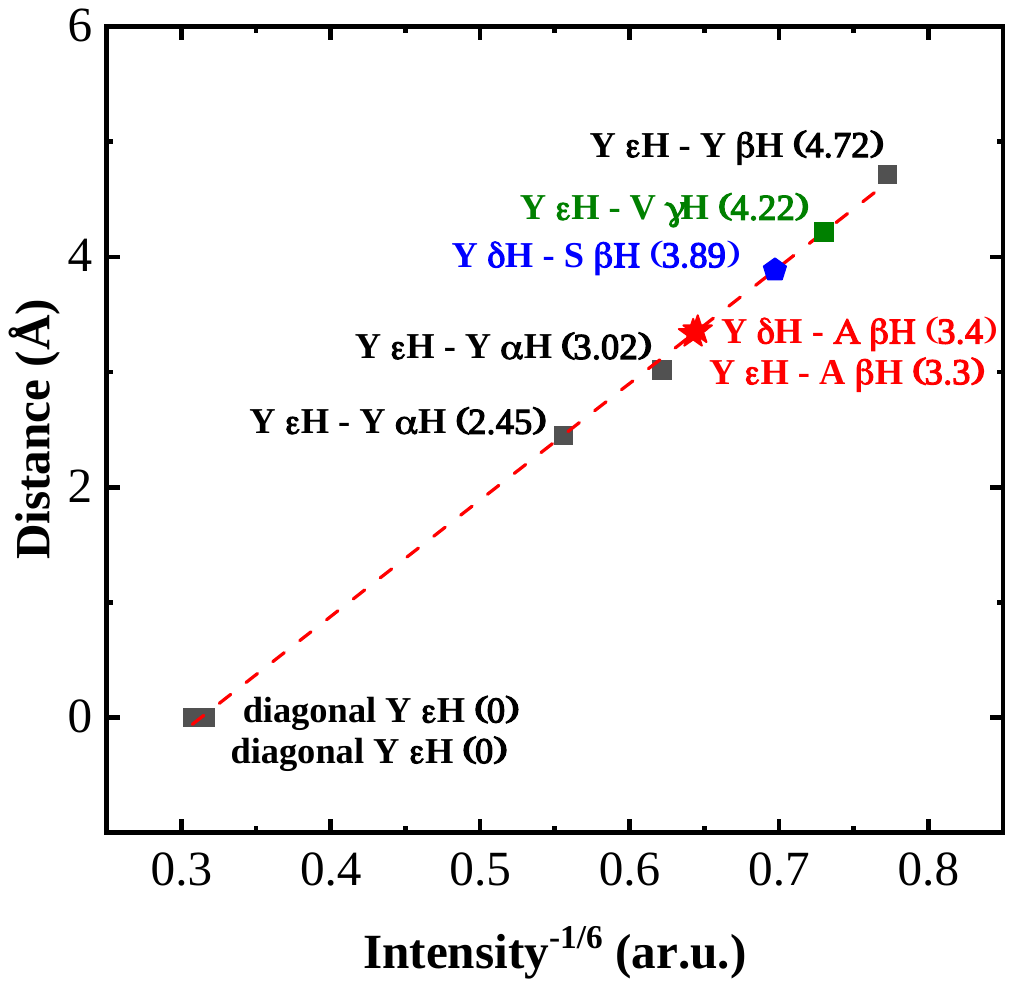

Fig. S11.

**Analysis and interpretation of NOESY data.** Distance against intensity ^-1/6^ for the assigned shifts and sticks representation of a Y residue showing estimated intra-residue distances used as internal calibration and estimated experimental values according to the internal calibration.

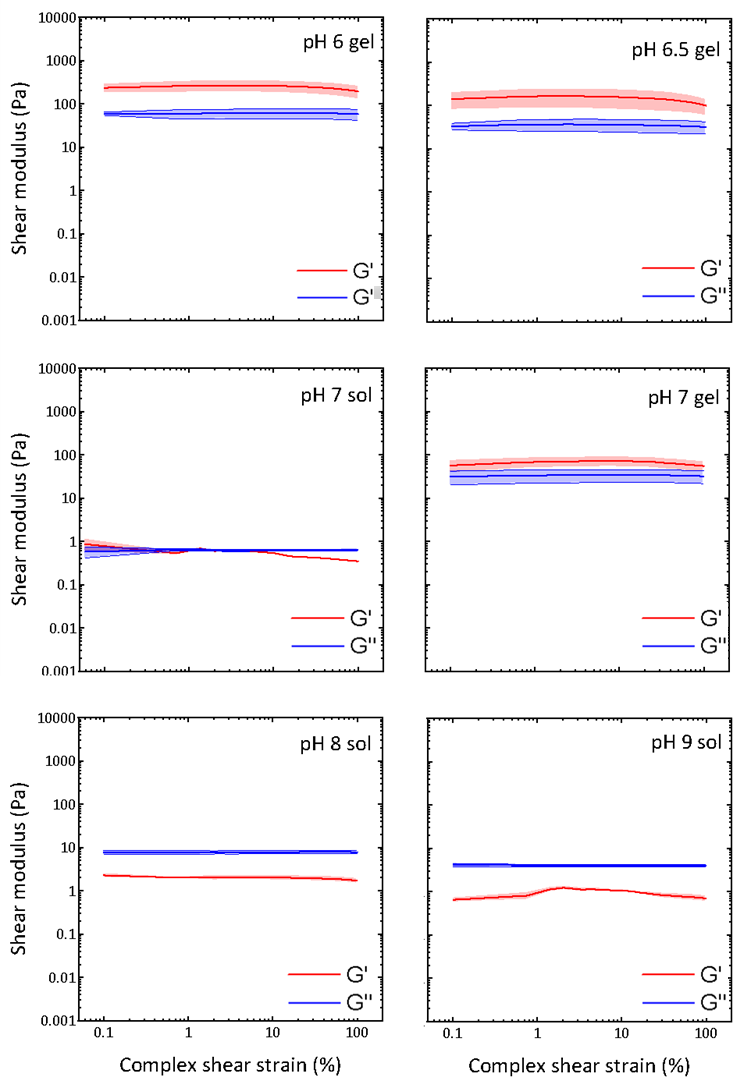

Fig. S12.

**Constant frquency rheology characterisation of NLSF.** Shear rheology oscillatory experiment where the complex strain was varied between 0.1 and 100 % at a constant frequency (1 Hz) for the samples of NLSF buffered at different pH (indicated within the graphs). Curves are shown as averages of 5 replicates, with the standard error shown as the shadowed area around the solid line. Experiments were intended to determine the linear viscoelastic limit of the materials.

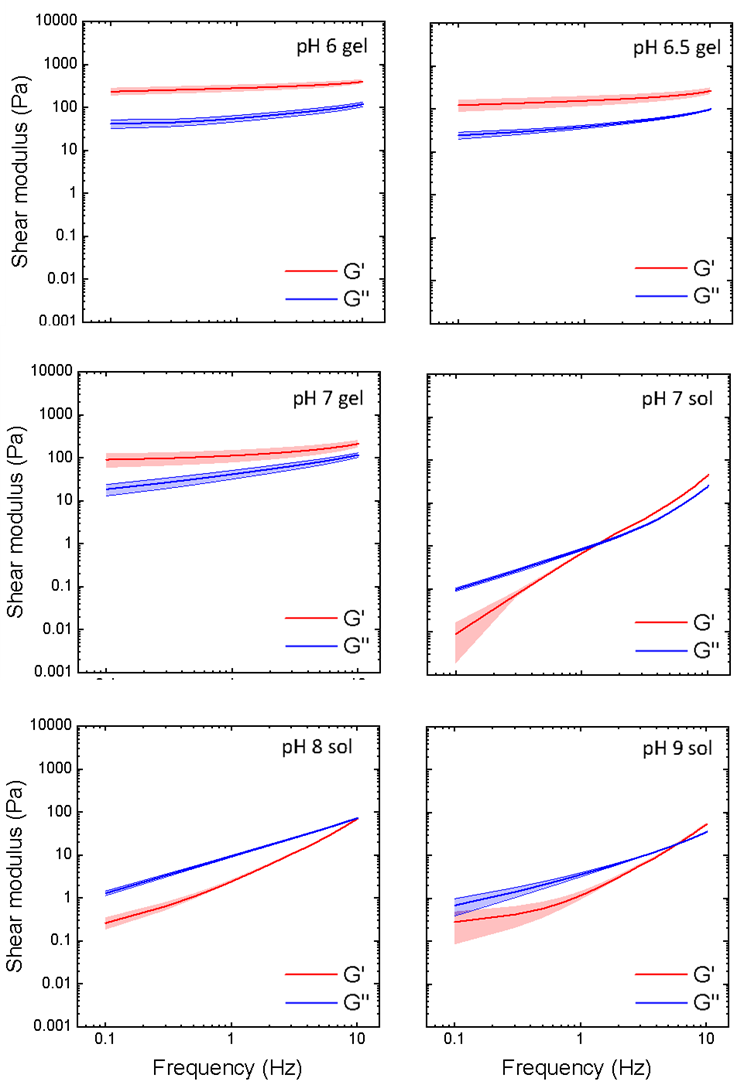

Fig. S13.

**Constant strain rheology characterisation of NLSF.** Shear rheology oscillatory experiment where the frequency was varied between 0.1 and 10 Hz at a constant strain (2 %) for the samples of NLSF buffered at different pH (indicated within the graphs). Curves are shown as averages of 5 replicates, with the standard error shown as the shadowed area around the solid line. Experiment shows the clear sol-gel transition suffered by the material as the pH is lowered below 7.

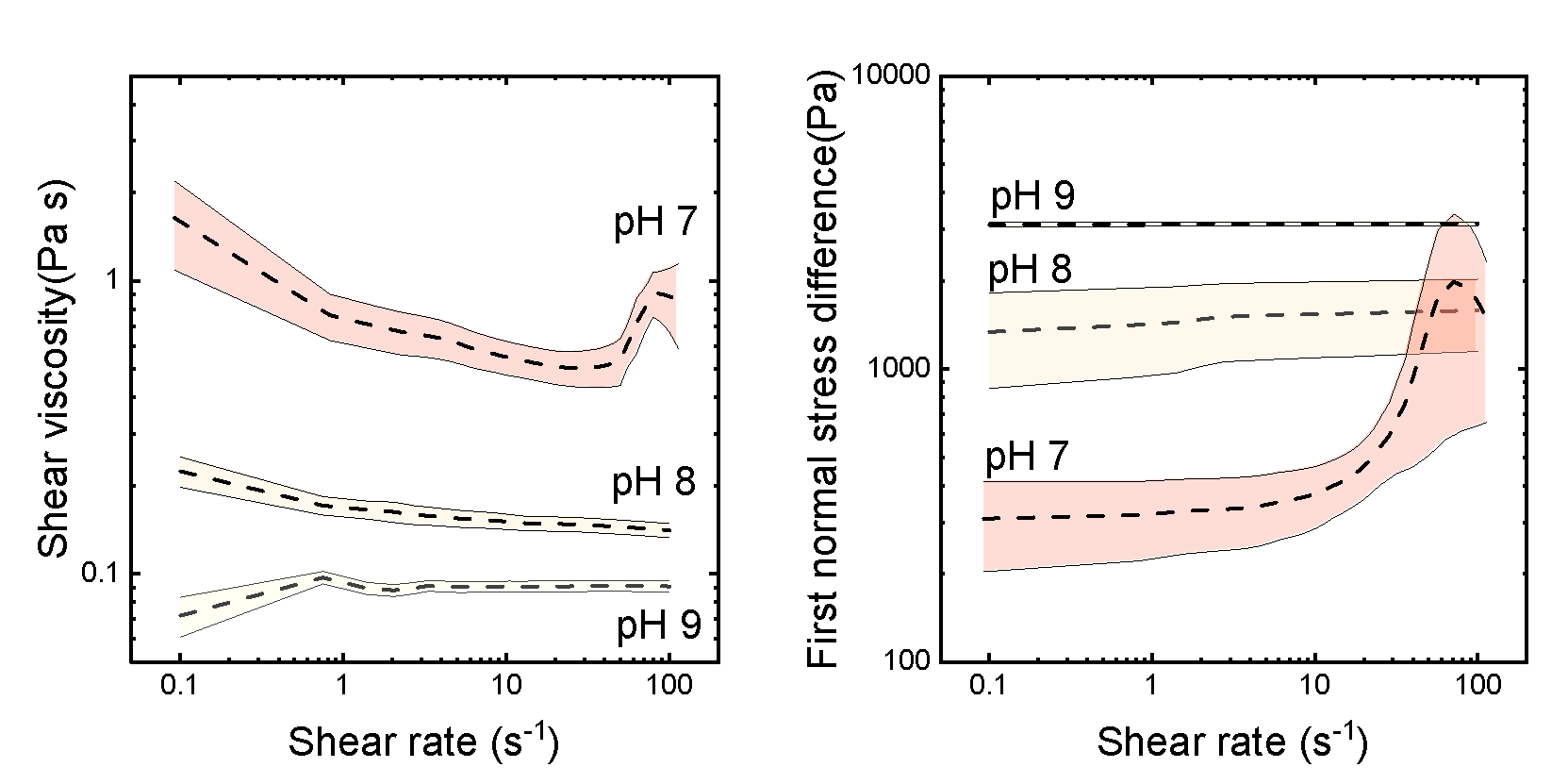

Fig. S14.

**Shear viscosity against shear rate for liquid samples of NLSF at different pH.** Samples at pH (7, 8 and 9) and (left) and first normal force difference vs shear rate of same samples. Curves showed averages (dashed black lines) and standard deviation (shadowed area around average), N = 5.

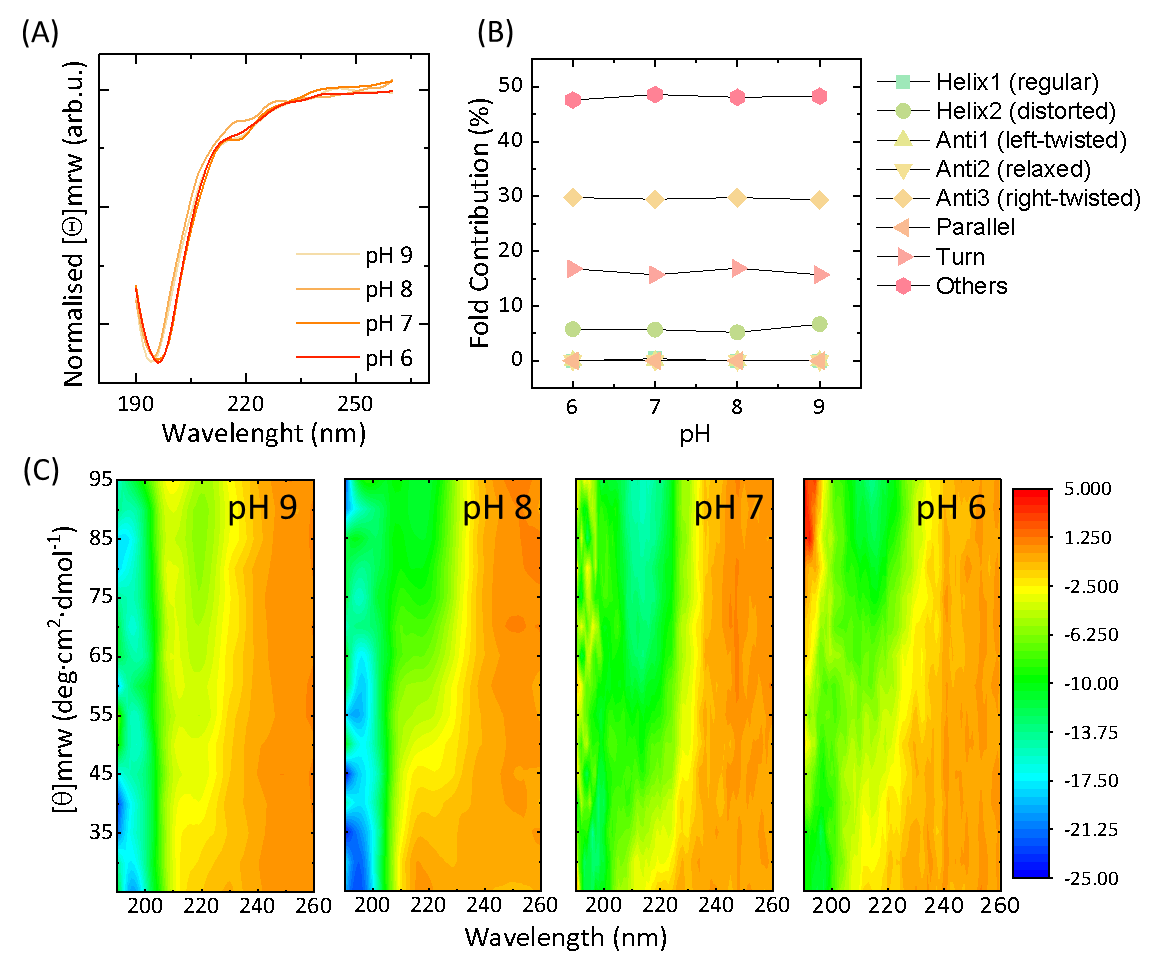

Fig. S15.

**Circular dichroism analysis of NLSF samples.** Overlapped normalized Ellipticity values of the buffered samples obtained by far-UV CD spectroscopy (A). Bestsel deconvolution of the curves showing estimated structural contributions to the ellipticity signal (B). Series temperature scans of the buffered samples in the far-UV region, where the vertical axis is the temperature, horizontal axis the wavelength, and the far-right the out-of-plane scale with red being the maximum range and blue minimum (C).

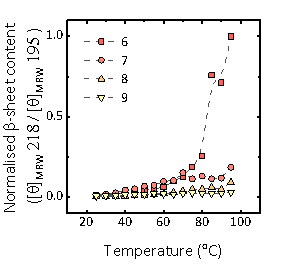

Fig. S16.

**Normalized β-sheet content as a function of temperature for all buffered samples.** The β-sheet content was estimated as the ratio between the ellipticity at 218 and 195 nm, normalized against the maximum obtained value for the data set.

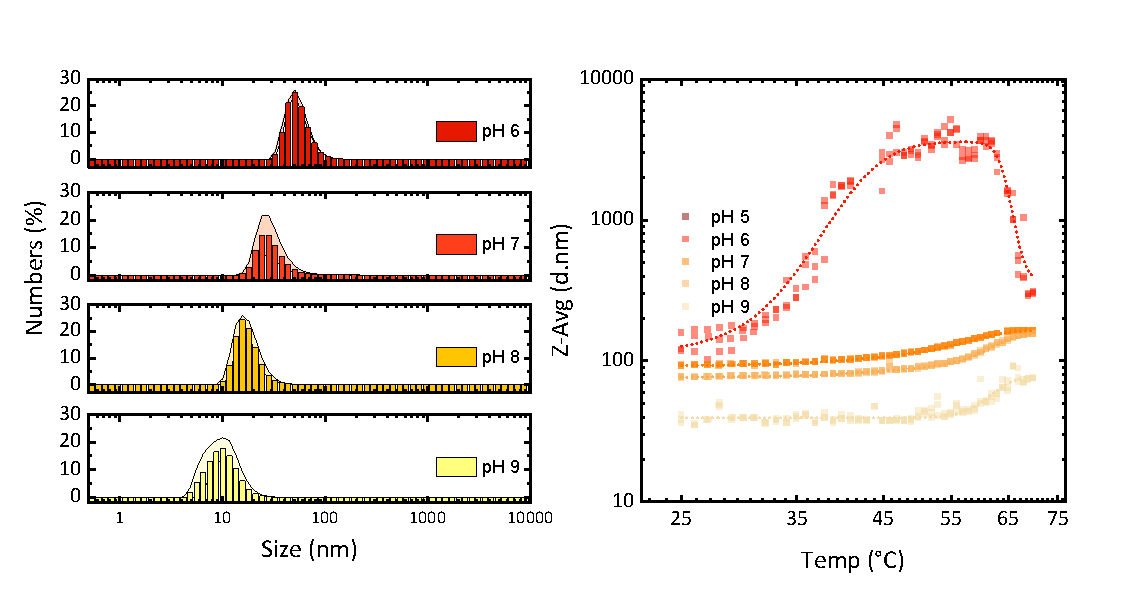

Fig. S17.

**Dynamic light scattering, DLS, analysis.** DLS size distribution of NLSF obtained at different pH values plotted as the Numbers (%) against the size in a lineal vs log scale (left). Values of the Z-average size for the buffered NLSF samples as a function of temperature (right).

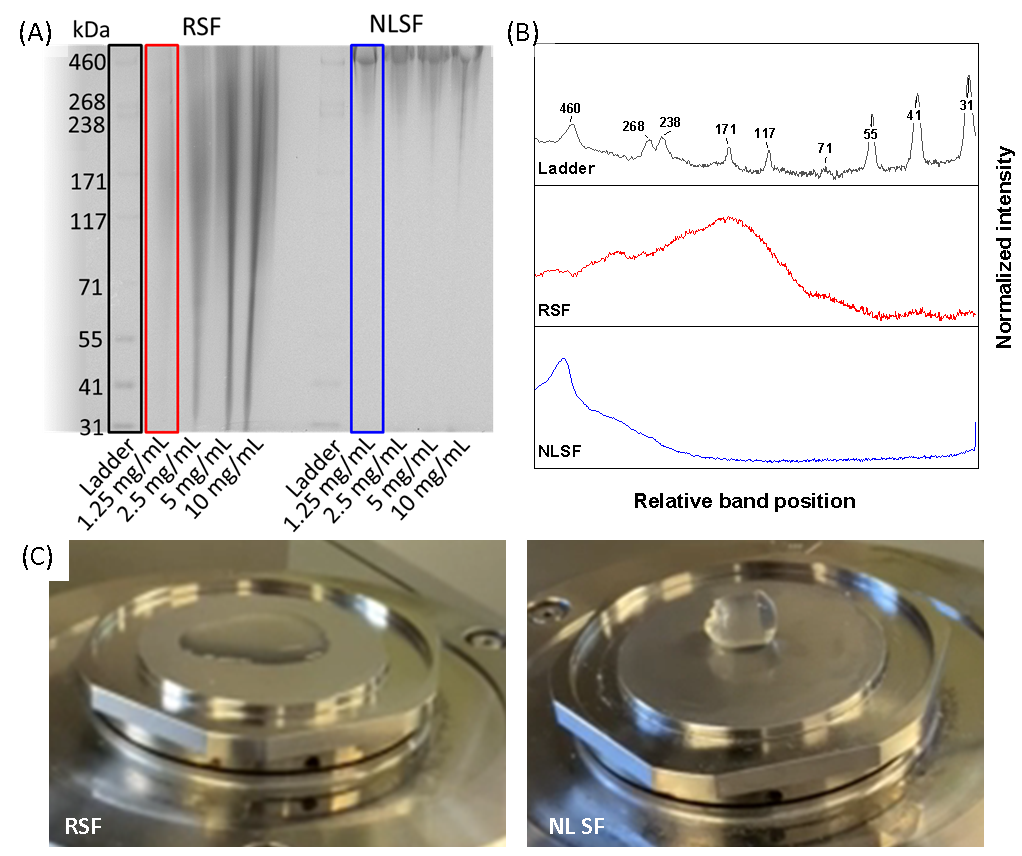

Fig. S18.

**Comparison of RSF and NLSF materials.** Uncropped SDS-PAGE gel of RSF and NLSF at different protein concentrations with reference ladder next to them (A). profiles extracted from the gels image from the selected bands in rectangles (B). Photos of samples loaded onto a rheometer geometry of unbuffered RSF and NLSF indicated (C).

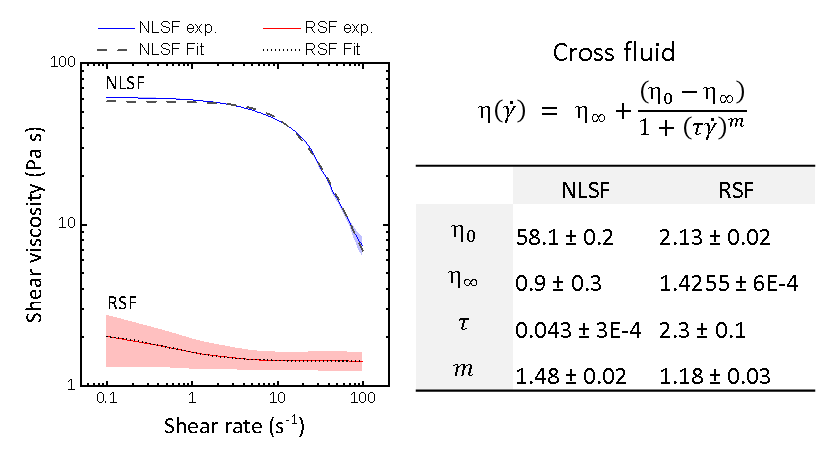

Fig. S19.

**Viscometry analysis of silk ion LiBr solutions.** Flow curves obtained for NLSf and RSF in their LiBr solutions plotted as shear viscosity against shear rate in a log-log scale (left). Solid lines represent experimental data, and dotted or dashed lines represent fits to a Cross fluid model, with constitutive equation shown on the right, and the table below it shows the fitting parameters.

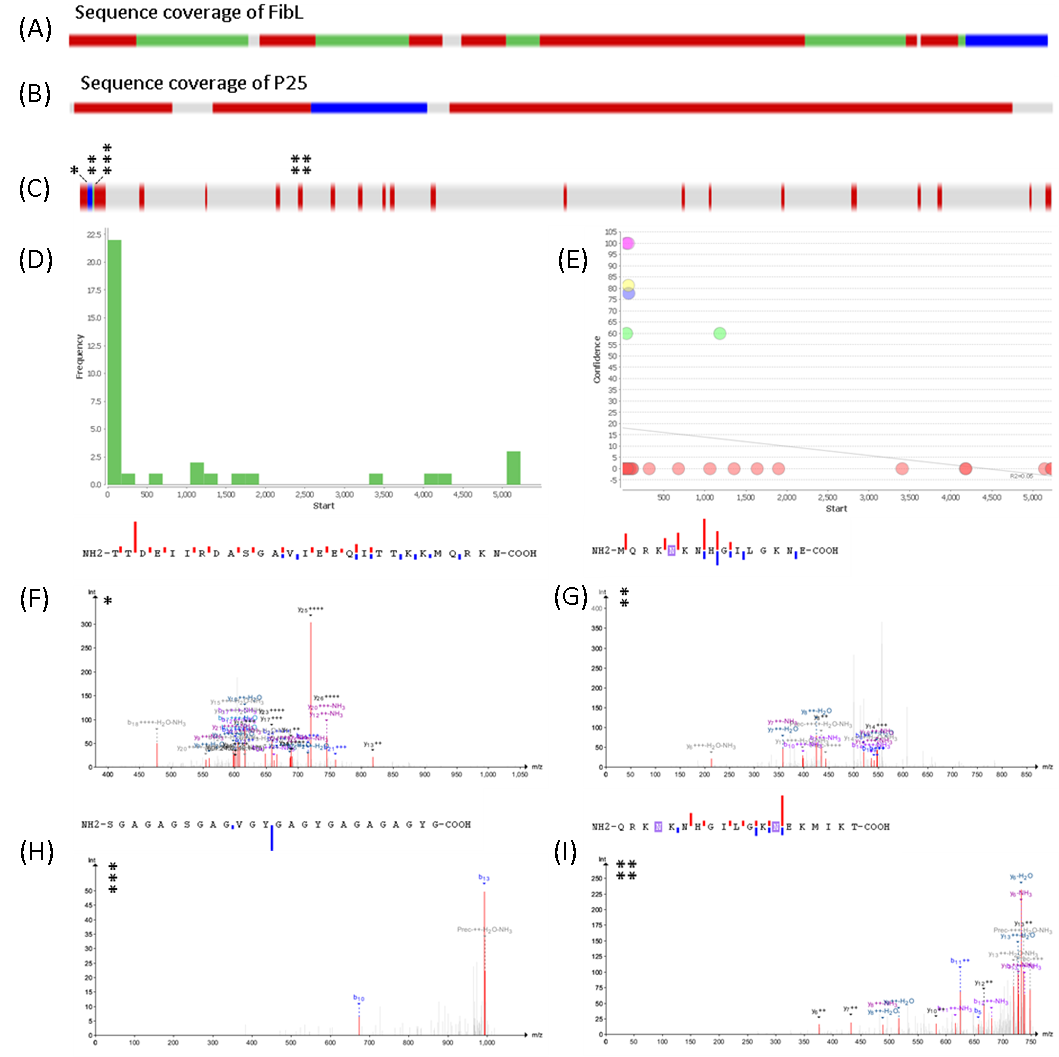

Fig. S20.

**Results from the proteomics analysis.** Cartoons representing the sequence coverage of FibL (A) P25 (B)) and FibH (C). Plot representing the frequency distribution of peptides against residue position of the covered peptides from FibH (D). Plot representing the confidence of found peptides against their residue position (E). Series of example mass spectrograms obtained for the labelled peptides in FibH (F-I). Within the cartoons, the red section represents low confidence assignments, and red and blue represent high confidence assignments.

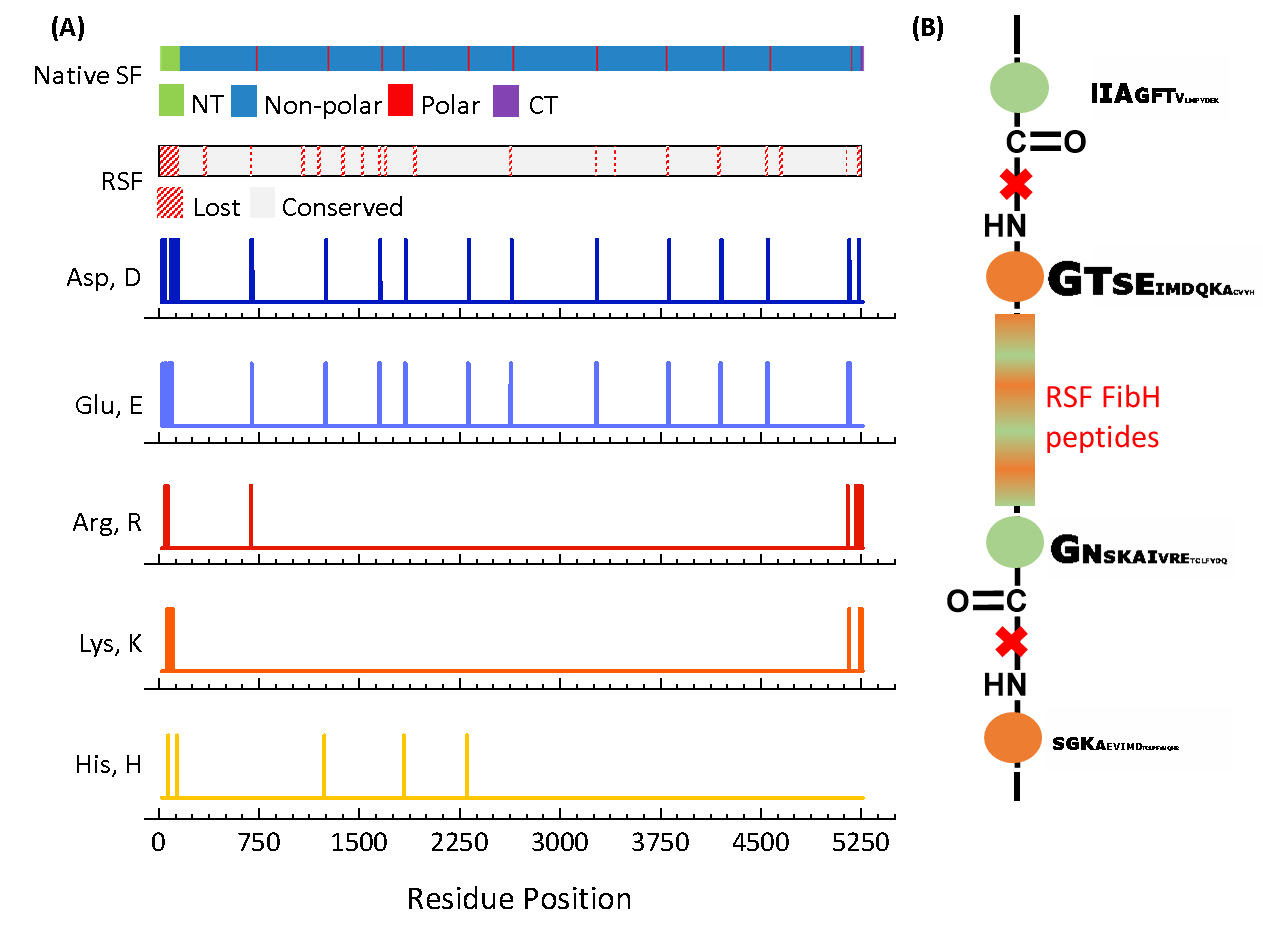

Fig. S21.

**Regeneration induced hydrolysis on fibroin**. Carton showing the multidomain architecture of the primary structure of FibH (top), followed by an aligned cartoon representing the total sequence coverage from the hydrolysate peptides recovered from RSF. Below, different plots show several indicated residues' positions along the sequence of FibH (A). Scheme representing the recovered peptides from RSF, with logo representations of the observed residues at terminal positions of the detected peptides, and those expected from the retained chains (B)

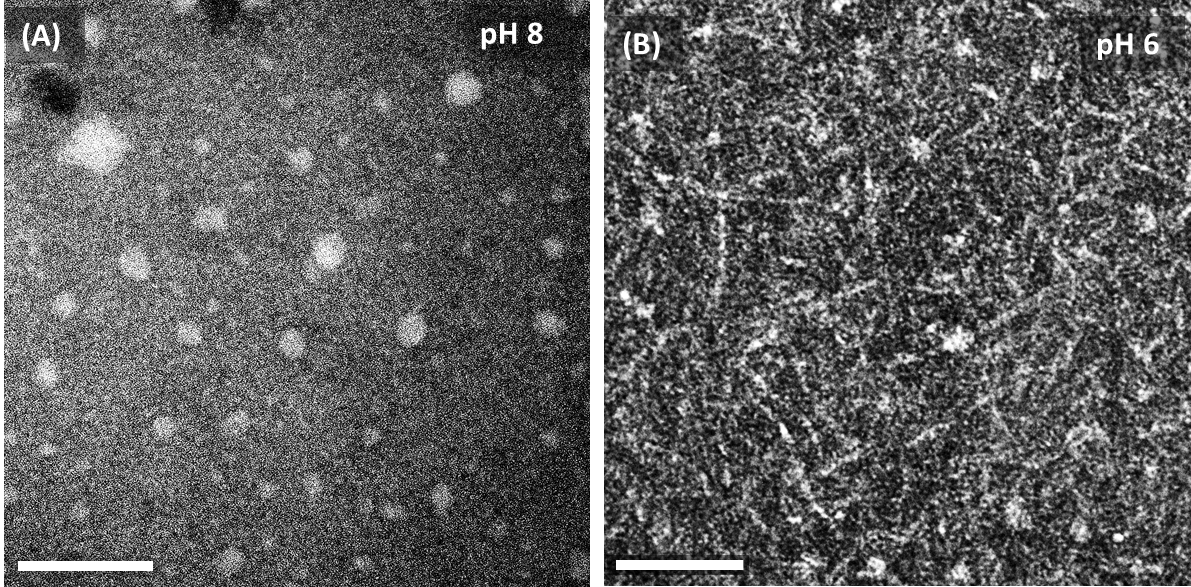

Fig. S22.

**TEM analysis of morphological changes in silk fibroin.** Negatively stained TEM images of globular units observed at pH 8 (A), and elongated fibrillar structures observed at pH 6 (B). Scale bars represent 100 nm in both cases.

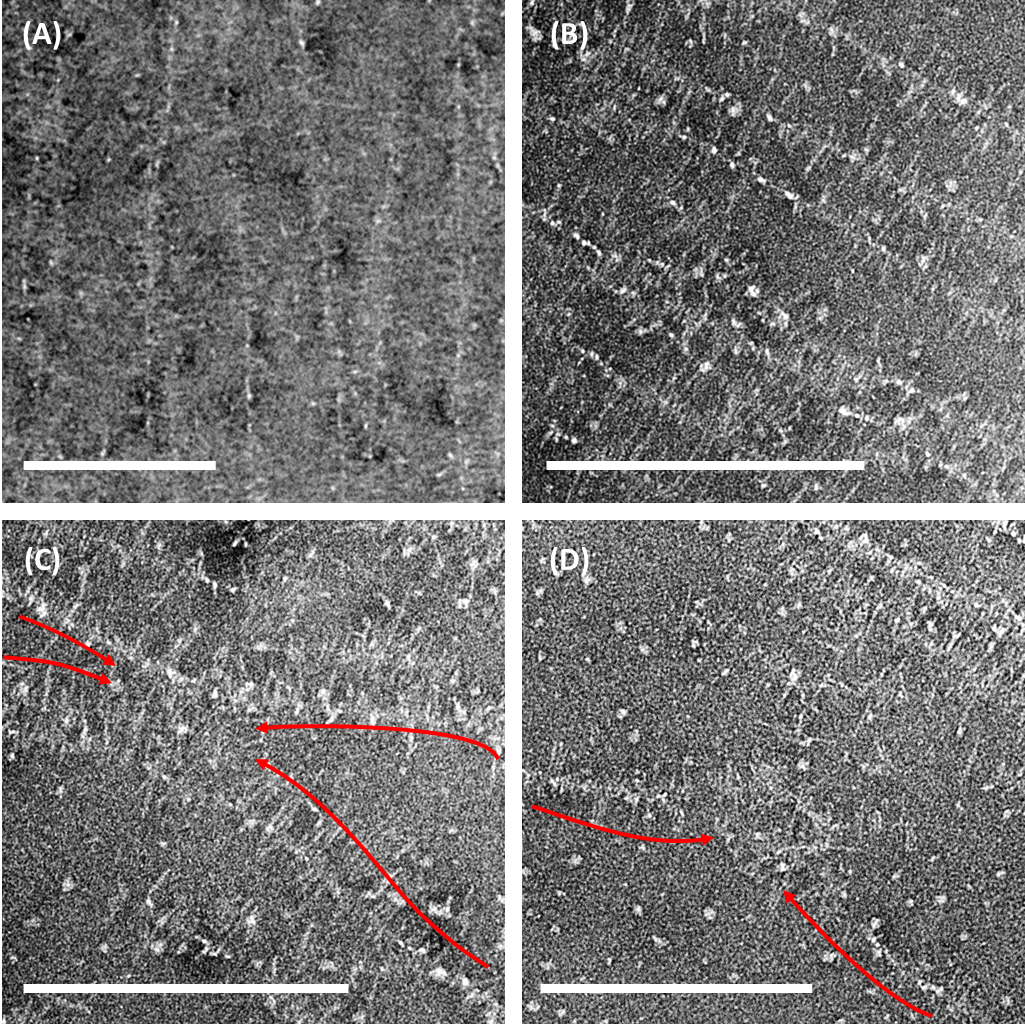

Fig. S23.

**TEM analysis of supramolecualr assemblies.** Negatively stained TEM images of intact supramolecular brush-like fibres (A, B) and illustrative images showing possible lateral interactions (C) and breakage of the structure (D), with arrows added as guides. Scale bars are 500 nm in (A-D).

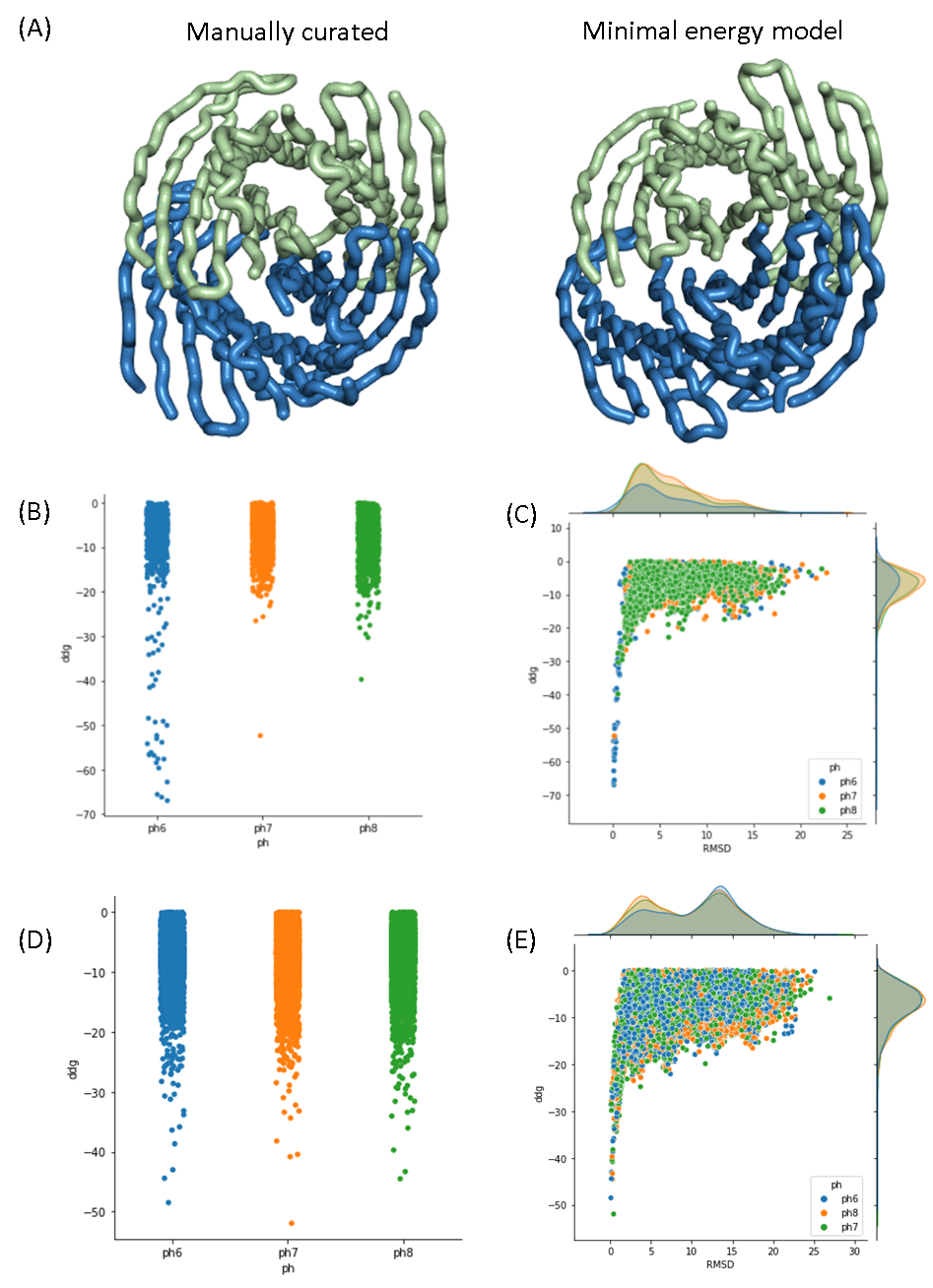

Fig. S24.

**N-terminal domain oligomerisation.** Ribbons representation of two stacked tetrameric units showing the manually curated model (left) and the equivalent relaxed and docked minimal energy model (right) as observed from the top (top).

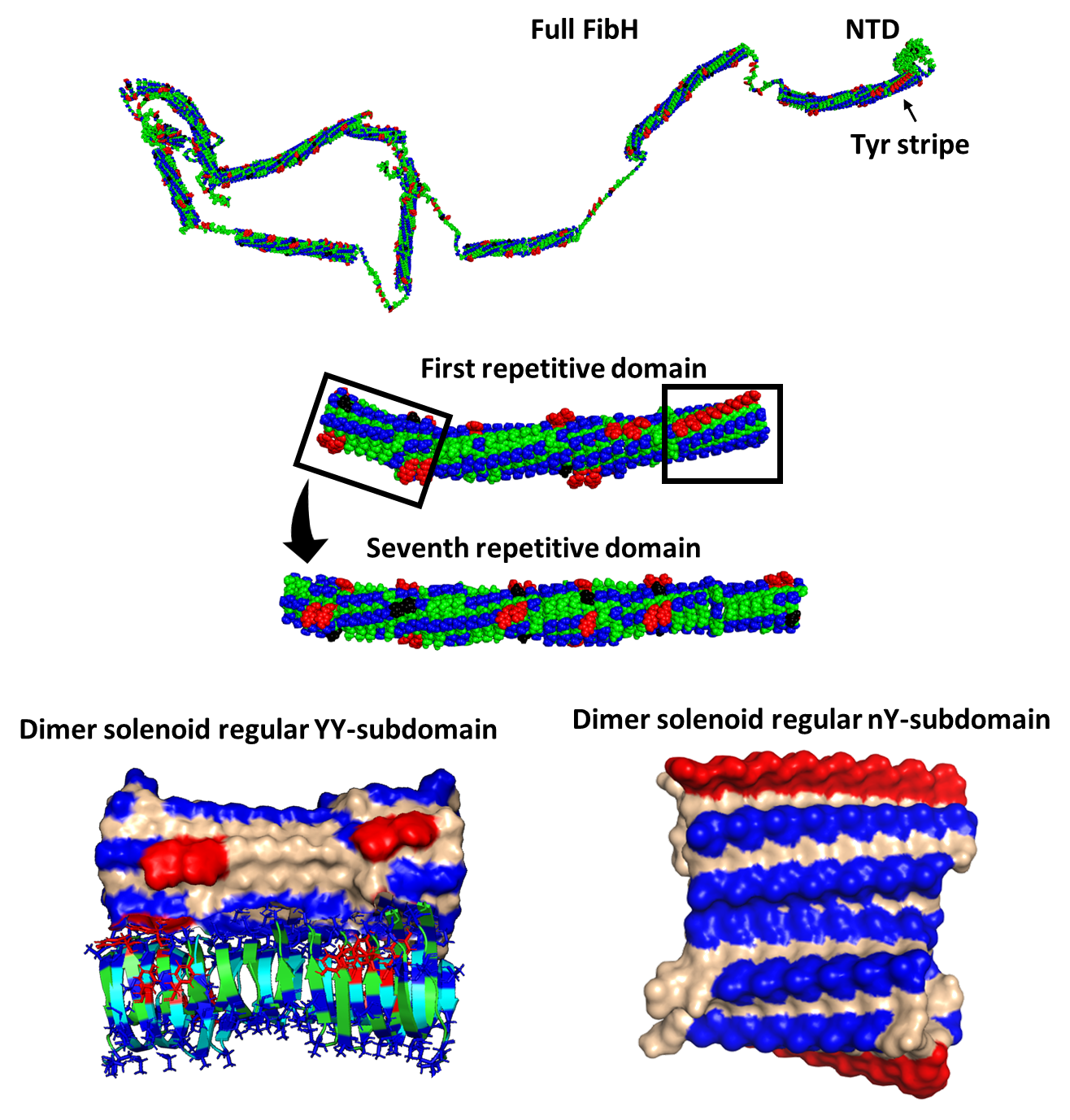

Fig. S25.

**The role of topology and Ala residues in driving lateral interactions**. Complete proposed FibH surface model coloured to highlight Ala (blue), Y (red) and Val (black) from all other residues (green) shown at the top. At the centre of the composite image, the detail of the first and sixth repetitive domain is shown, highlighting the detected subdomains and the difference in the spatial distribution of Y residues. At the bottom, two laterally docked models of the two different subdomains, showing binding at Ala interfaces with avoidance of Y residues; colour wheat or green represent all other residue but Ala and Y.

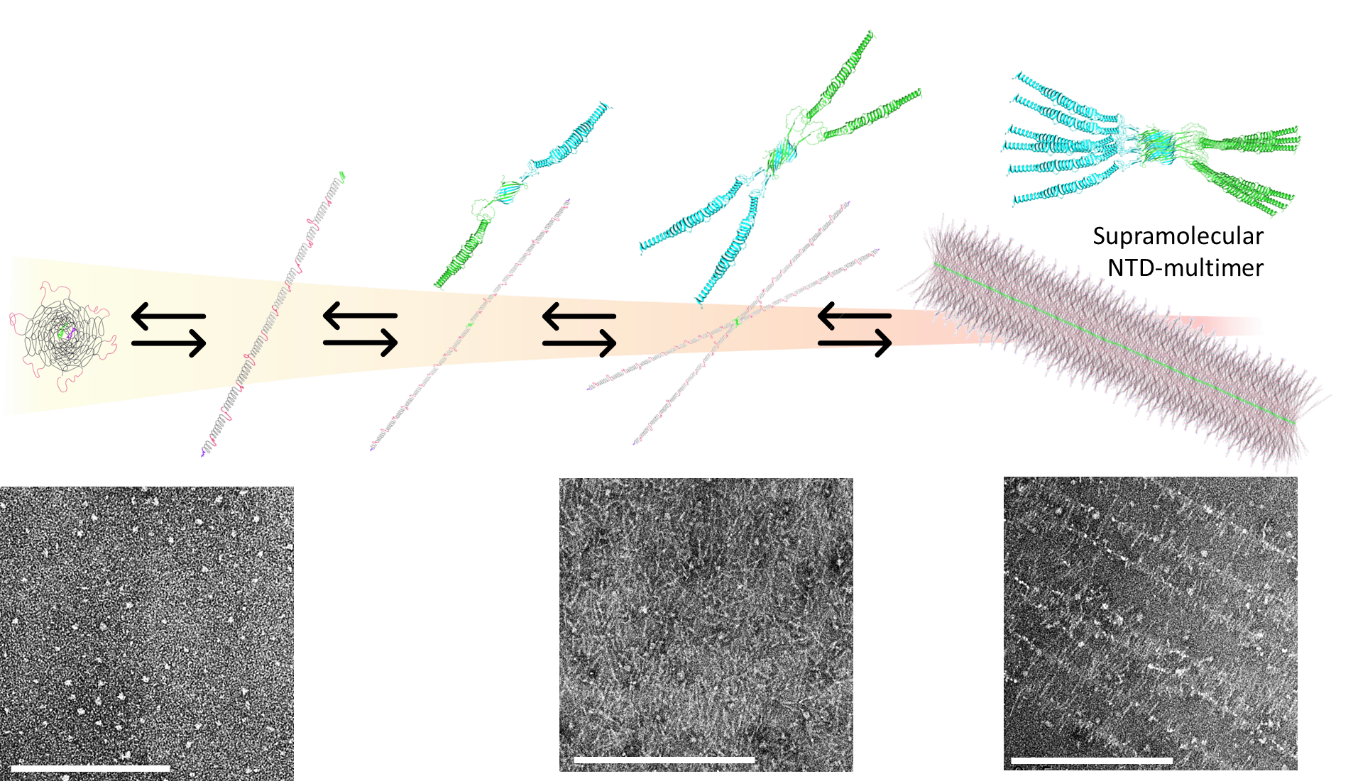

Fig. S26.

**Proposed NTD-driven self-assembly process**: FibH, as a multidomain solenoidal molecule exists, takes a globular appearance with a diameter of about 20 nm at pH values above 7. At pH 7, the linkers are partially protonated, and the protein can extend and start participating in lower-order oligomer species. As pH continues to drop, assembly continues into the supramolecular bottlebrush fibres. PDB models represent only the NTD with the first repetitive domain, whereas drawings represent, schematically, the complete molecule. The background image's colour and shape allude to the pH gradient (from 8 to 6) and the anterior section of the silk gland (ASG), respectively. Negatively stained TEM images shown below are intended to show snapshots of the assembly process, and scale bars represent 500 nm.

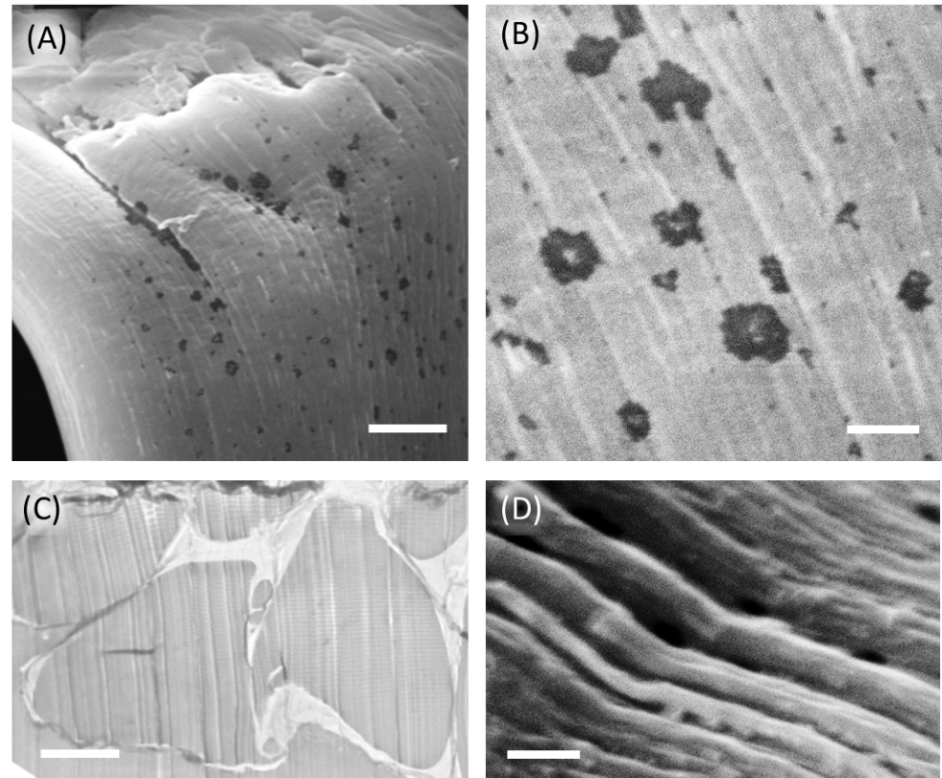

Fig. S27.

**Microscopy analysis of natural silk fibre.** SED-SEM image of a sheared degummed fibre (A). SEM detail off the same fibre showing nanofibrillar texture and void formation during shearing (B). TEM image of a microtomed section of an embedded raw fibre showing twin fibroin fibres coated by a less electron-dense sericin layer (C). SEM image showing further detail of the nanofibrillar texture on a tilted specimen (D). Scale bars are 2 µm in (A) and 500 nm (B-D).

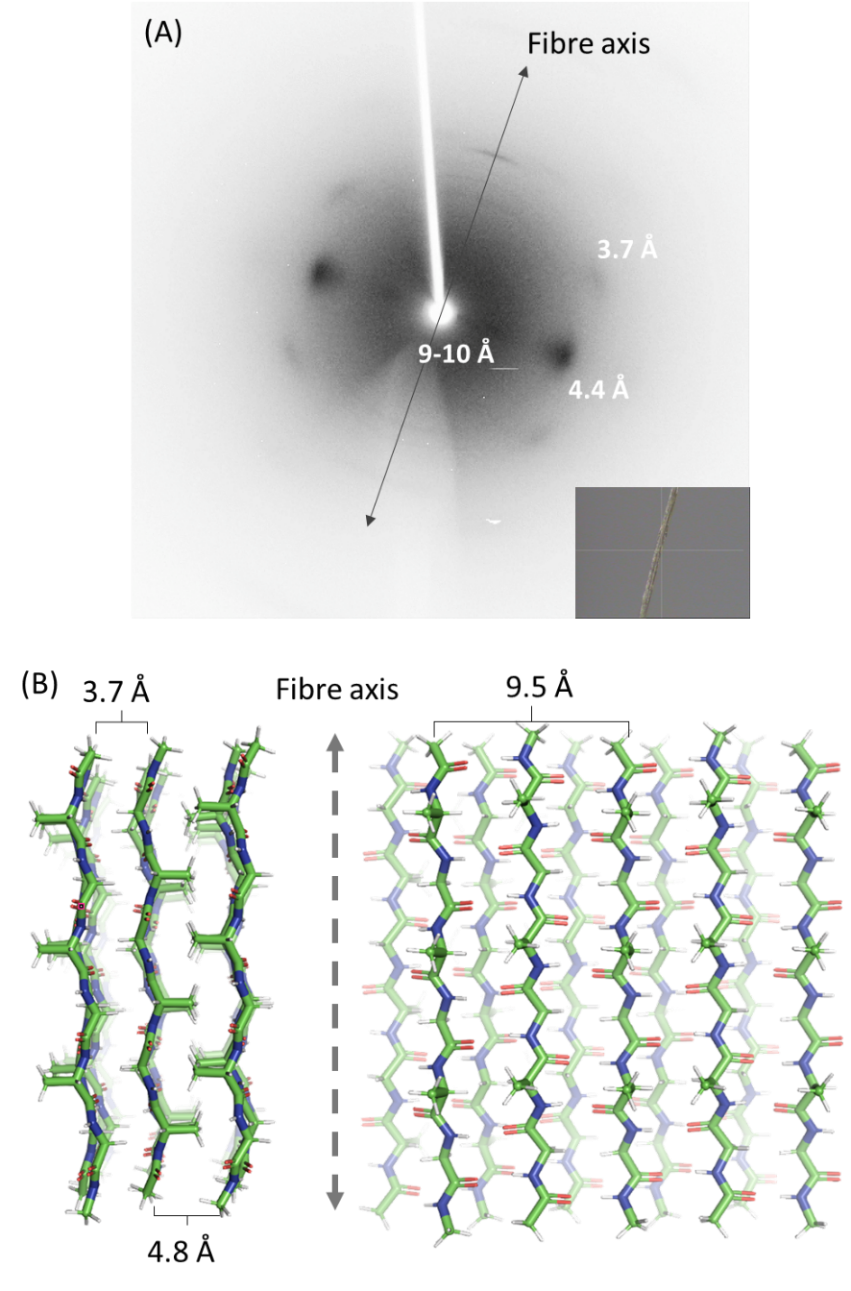

Fig. S28.

**Fibre X-ray diffraction of natural silk fibre.** 2D X-ray diffraction pattern of a raw silk fibroin fibre with assigned distances (A) atomic model of the consensus Silk-II structure, showing the polar antiparallel β-sheet with alternating A-A and G-G interfaces (B). The model used was extracted from the model archive website (DOI: [10.5452/ma-cs24y](https://dx.doi.org/10.5452/ma-cs24y)), proposed before (*85*).

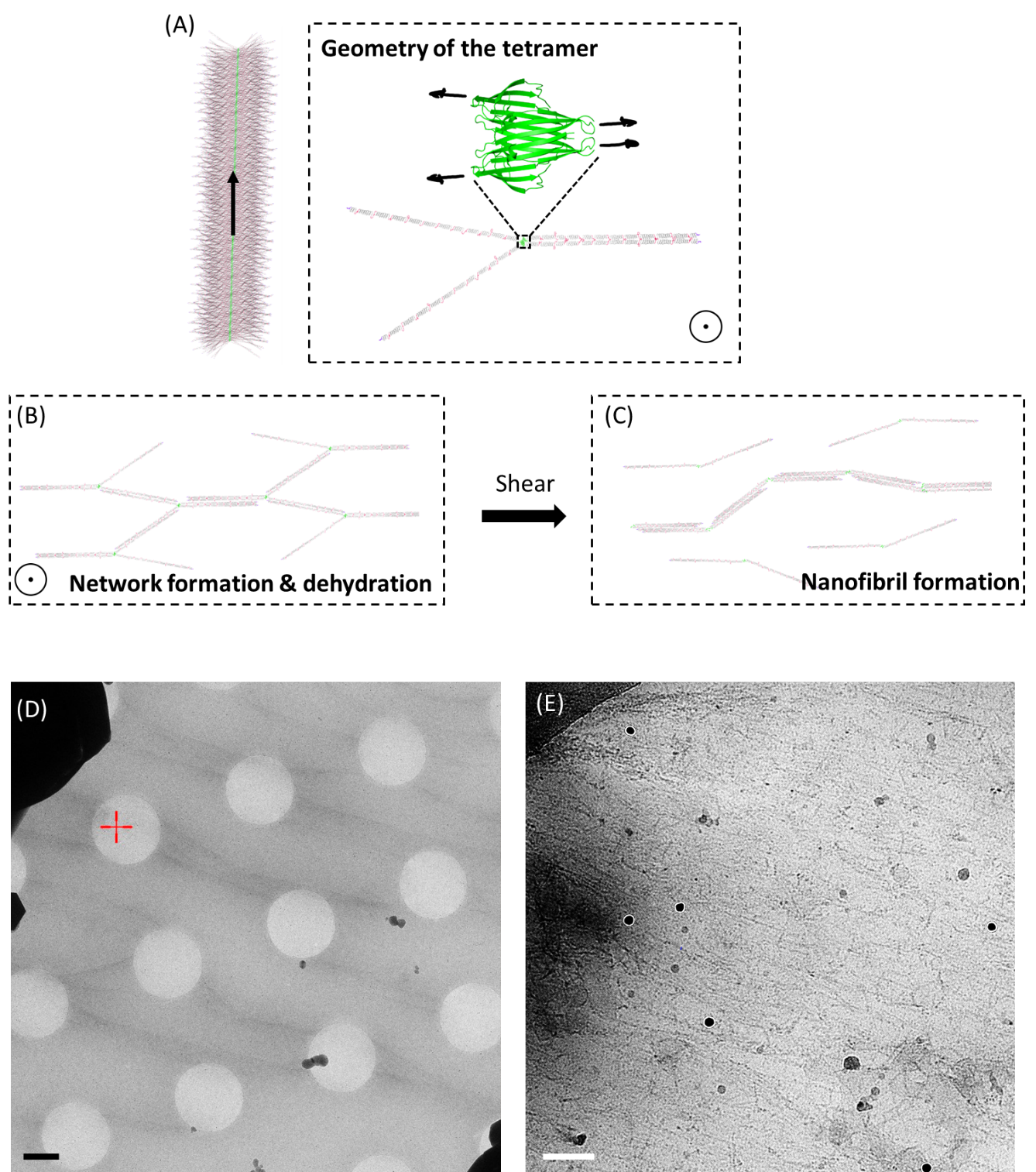

Fig. S29.

**Proposed network formation is driven by lateral interaction of solenoid units:** Structure of a tetrameric unit within the supramolecular complex, showing the direction towards c-termini where the first repetitive domain would be located and the possible initiation intra-unit lateral interactions (A). Network of supramolecular fibres stabilized by lateral interactions of solenoids (B). Breakage of the NTD interactions and transformation to a flow-aligned solenoid network formation (C). Cryo-TEM image obtained from an apparent proto fibre at low magnification (D), and at high magnification (E), showing what is believed to be evidence of the pre-extended solenoid network.

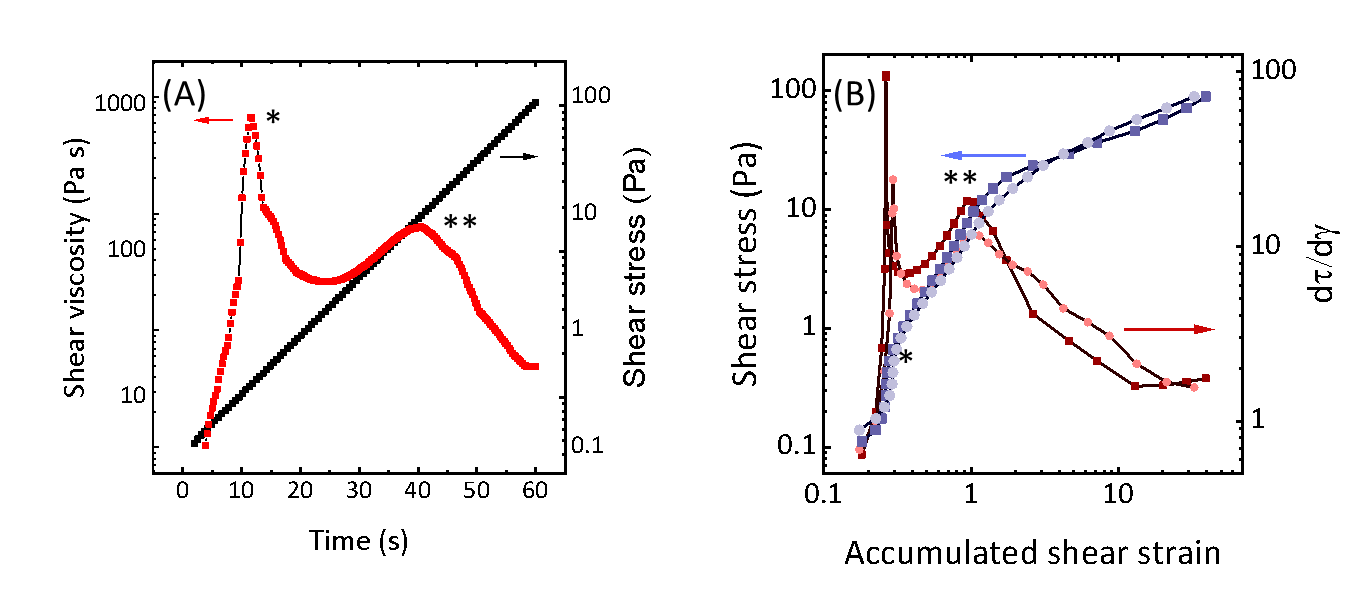

Fig. S30.

**Evidence of two yield stresses.** Shear yield stress experiments conducted on a pH 7 viscoelastic sample: Shear viscosity and shear stress against time in a log-lineal scale, showing two yield points shown as maxima in viscosity (A). Shear stress and its derivative against accumulated shear strain for two independent samples showing a first sharp and a second broad yield (B).

Table S1.

Reflections obtained from the oriented film diffraction pattern show typical Silk-I spacing.

| Experimental results | | | | |
| --- | --- | --- | --- | --- |
| Equatorial | |  | **Meridional** | |
| d-Spacings (Å) | Intensity |  | d-Spacings (Å) | Intensity |
| 7.35 | 226 |  | 5.21 | 169 |
| 5.12 | 132 |  | 3.50 | 111 |
| 3.70 | 170 |  | 3.15 | 104 |
| 2.56 | 62 |  | 2.04 | 42 |
| 2.06 | 33 |  |  |  |

Table S2.

**NMR chemical shifts.** Assigned chemical shifts for simplified motifs compared to those found in the literature (*43*)**.**

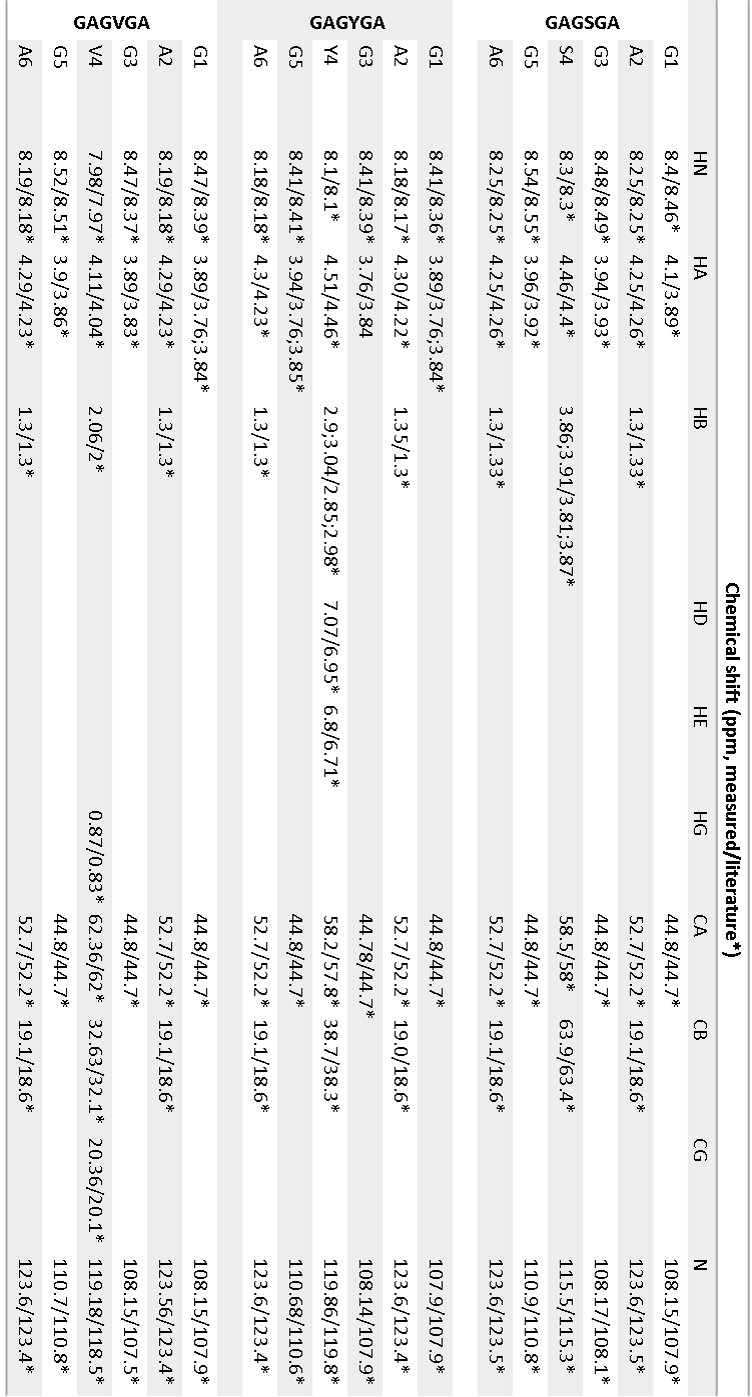

Table S3.

**Predicted chemical shifts for simplified motifs within the sixth repetitive domain**. Chemical shifts were predicted using the webserver from SHIFTX2.

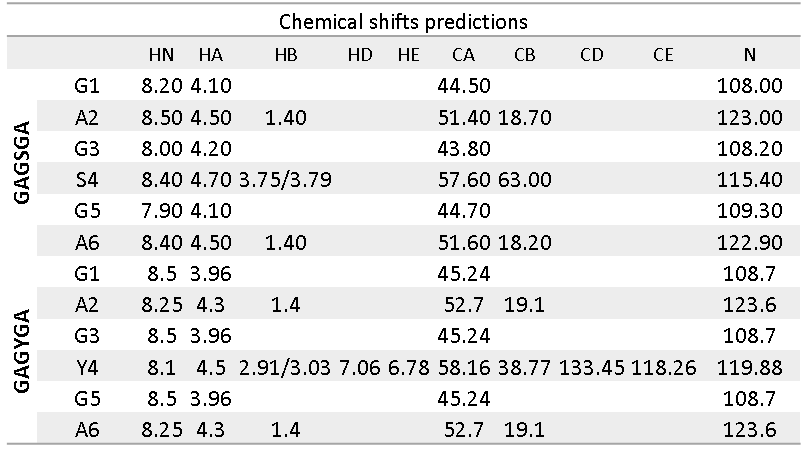

Table S4.

Table with the logarithmic shifts used to calculate the master curves

| Sol master curve | | | Gel master curve | |
| --- | --- | --- | --- | --- |
| pH | Log(a_pH_), Freq. | *****Log(a_pH_), G | *pH* | Log(a_pH_), Freq. |
| 7 | 0 | 1.83 | 7 | 0 |
| 8 | -0.86 | 0 | 6.5 | 1.23 |
|  |  |  | 6 | 3.14 |

*****Given the suspected lower concentration of the sol fraction at this pH, and knowing that like viscosity, the moduli follow a power-law relationship with concentration, the moduli were scaled to match that of pH 8, however leaving the frequency domain unaffected.
